# Vascular dimorphism ensured by regulated proteoglycan dynamics favors rapid umbilical artery closure at birth

**DOI:** 10.1101/2020.07.02.184978

**Authors:** Sumeda Nandadasa, Jason M. Szafron, Vai Pathak, Sae-Il Murtada, Caroline M. Kraft, Anna O’Donnell, Christian Norvik, Clare Hughes, Bruce Caterson, Miriam S. Domowicz, Nancy B. Schwartz, Karin Tran-Lundmark, Martina Veigl, David Sedwick, Elliot H. Philipson, Jay D. Humphrey, Suneel S. Apte

## Abstract

The umbilical artery lumen occludes rapidly at birth, preventing blood loss, whereas the umbilical vein remains patent, providing the newborn with a placental infusion. Here, we identify differential arterial-venous proteoglycan dynamics as a determinant of these contrasting vascular responses. We show that the umbilical artery, unlike the vein, has an inner layer enriched in the hydrated proteoglycan aggrecan, external to which lie contraction-primed smooth muscle cells (SMC). At birth, SMC contraction drives inner layer buckling and centripetal displacement to occlude the arterial lumen, a mechanism elicited by biomechanical and computational analysis. Vascular dimorphism arises from spatially regulated proteoglycan expression and breakdown in umbilical vessels. Mice lacking aggrecan or the metalloprotease ADAMTS1, which degrades proteoglycans, demonstrated their opposing roles in umbilical cord arterial-venous dimorphism and contrasting effects on SMC differentiation. Umbilical vessel dimorphism is conserved in mammals, suggesting that their differential proteoglycan dynamics were a positive selection step in mammalian evolution.

## INTRODUCTION

The umbilical cord, typically containing two arteries and one vein in humans, is a crucial fetal structure in placental mammals. Umbilical arteries carry fetal blood to the placental vascular bed whereas the umbilical vein returns oxygenated blood to the fetus. Fetal respiration at birth renders the maternal oxygen supply redundant. Umbilical arteries commence closure rapidly after delivery of the newborn whereas the veins remain open longer. The cord is routinely clamped following delivery and divided between the clamps in modern obstetric practice and timing of cord clamping after birth, whether early or late, is extensively debated (1, 2). A recent recommendation suggested clamping 30-60 seconds after birth to facilitate the placental transfusion (3). Although the necessity of clamping is rarely questioned, it appears to be a modern practice (4). Cord clamping is not practiced in domesticated animals and wild animals, yet all mammalian species have survived. We hypothesized that intrinsic design characteristics of mammalian umbilical arteries may prevent blood loss at birth without clamping.

Prior work has noted that umbilical arteries have a bilaminar structure histologically (5), in addition to the absence of elastic lamellae, which endow large arteries with resilience to cyclic loading (6), but its underlying molecular mechanisms and relationship to arterial closure remained obscure. Here, we used a multi-disciplinary approach integrating a variety of morphologic approaches with mechanical testing, computational analysis and insights from mouse mutants to demonstrate the molecular and biomechanical basis for rapid umbilical artery closure. The findings emphasize the dual importance of extracellular matrix proteoglycans in regulation of cell differentiation and conferment of desirable tissue mechanical characteristics.

## RESULTS

### The umbilical artery has a bilaminar wall

Three-dimensional imaging of term human umbilical cords, using synchrotron-based phase contrast micro-CT (effective pixel size 1.63 x 1.63 μm^2^) (7) and histology identified a much thicker tunica media (TM) in the umbilical artery than the vein, with a visibly different structure (Fig. 1a-c, Sup. Fig. 1a-c, Sup. movies 1,2). Most umbilical arteries were occluded at birth independent of delivery method or cord region analyzed, whereas umbilical veins remained patent (Fig. 1b, Sup. Fig. 1b). Smooth muscle cell (SMC) markers showed similar staining intensity within inner and outer TM of the umbilical arteries and TM of vein (Sup. Fig. 1c,d).

**Figure 1.**
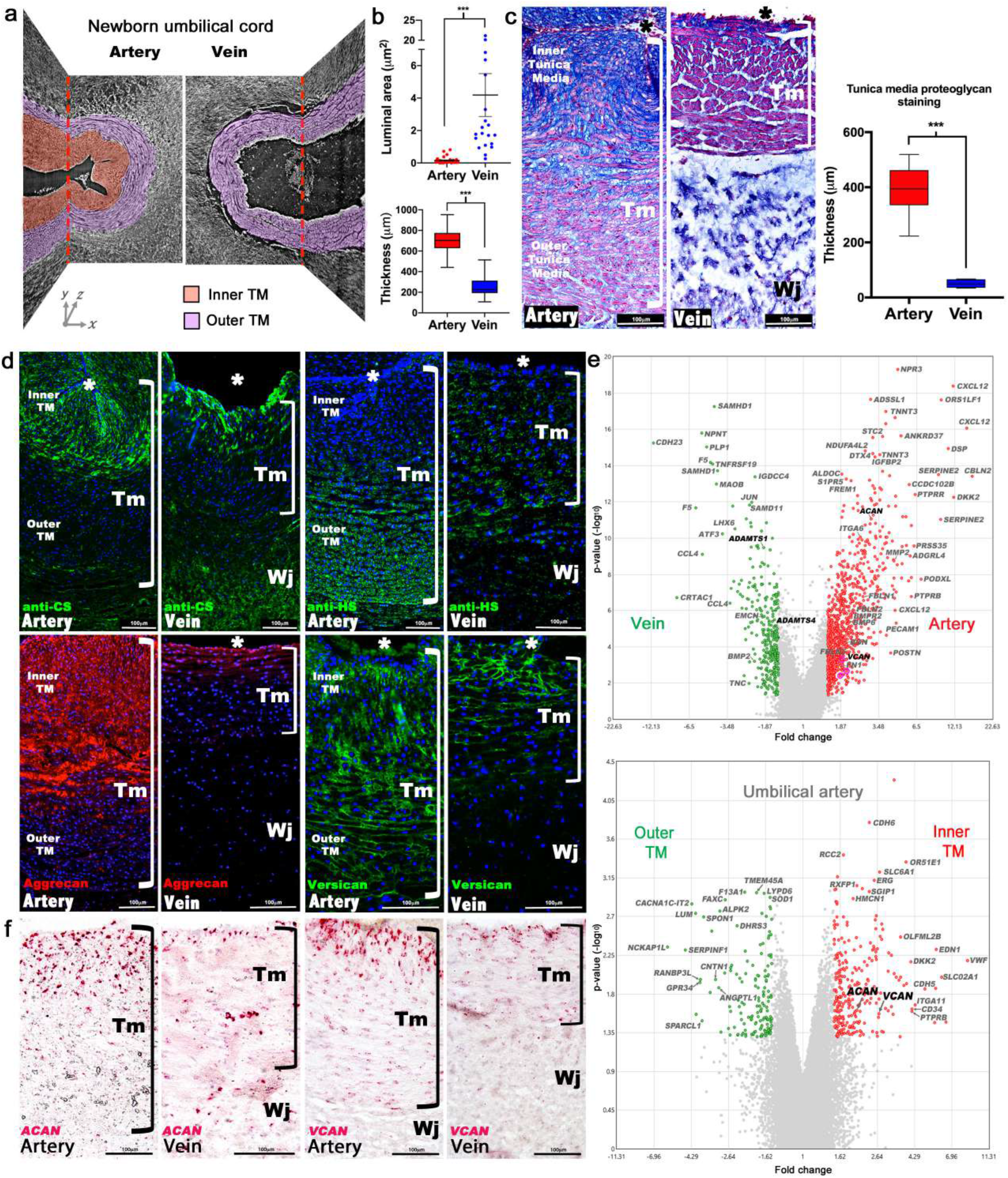
Dimorphism of the human umbilical artery and vein. **(a)** Synchrotron imaging of umbilical vessels at birth illustrates a bilayered arterial wall comprising an inner buckled tunica media (TM, red) and outer TM (purple) but no distinct inner layer in the vein. X-Y and Y-Z image planes are indicated by red dashed lines. **(b)** Quantitation of luminal area at birth shows that the umbilical arteries are occluded whereas the veins remain patent (top) and significantly thicker arterial wall (bottom) (n=20 cords, error bars indicate mean ± S.E.M., whiskers indicate minimum and maximum values. ***, p<0.001). **(c)** Alcian blue, eosin (pink) and nuclear fast red staining of umbilical vessel cross-sections shows a proteoglycan-rich (blue) inner TM in the umbilical artery but not the vein. Quantified staining intensity is shown on the right (n=6 umbilical cords, whiskers indicate minimum and maximum values, ***, p<0.001. (**d**) Chondroitin sulfate (CS), heparan sulfate (HS), aggrecan and versican immunofluorescence showing that CS staining corresponds with aggrecan and versican staining and alcian blue in (c). (**e**) Volcano plots illustrating differential gene expression between human umbilical artery (red) and vein (green) (top, n=4 umbilical arteries and veins) and differential gene expression between human umbilical artery inner (red) and the outer TM (green) (bottom, n=2 inner and outer TM). (**f**) In situ hybridization shows robust ACAN and VCAN expression in the inner artery TM and weak expression in the vein. * marks the vessel lumen. Brackets in **c,d,f** mark the TM. Wj, Wharton’s jelly. Scale bars = 100µm in **c,d,f**.

Alcian blue, which binds sulfated glycosaminoglycans (GAGs) intensely stained the inner TM but only the innermost 3-4 cell layers of the venous TM (Fig. 1c). SMCs in this GAG-rich region of the arteries were radially oriented and round, with nuclear-localized Sox9, a chondrogenic factor (Sup. Fig.1e) (8). The distribution of chondroitin sulfate (CS) coincided with Alcian blue staining (Fig. 1c-d), whereas heparan sulfate was more abundant in the outer TM cells (Fig. 1d), suggesting that the inner TM was enriched in CS-proteoglycans (CSPGs). RNA microarray data from matched human umbilical arteries and veins showed, among many differentially expressed genes (Fig. 1e, Sup. Fig. 2, Sup. Table 1), arterial prevalence of mRNAs encoding CSPGs with the most CS-chains, aggrecan and versican (Fig. 1d). Microarray analysis of the inner versus outer arterial TM identified stronger *ACAN* and *VCAN* expression in the inner TM, among other differences (Fig. 1d, Sup. Fig. 3, Sup. Table 2). RNA *in situ* hybridization (RNA-ISH) localized strong *ACAN* and *VCAN* expression in arterial inner TM SMC, and immunostaining showed versican and aggrecan in a similar distribution as alcian blue and anti-CS staining (Fig. 1c, d). Versican is a well-characterized vascular component (9), but the abundance of aggrecan, known as a cartilage/neural proteoglycan (10, 11) was unexpected.

**Table 1.** Model parameters fixed for all simulations of the umbilical artery, determined primarily from the biaxial biomechanical data and histological findings.

| Parameters | Description | Values |
| --- | --- | --- |
| $A, B, C$ | Unloaded inner, interface, outer radius | 173.65, 208.50, 231.73 $\mu\text{m}$ |
| $\lambda_z$ | Loaded axial stretch | 1.25 |
| $\mu_1, \mu_2$ | GAG/matrix shear modulus inner, outer layer | 3.2 kPa, 0.1 kPa |
| $c_1^1, c_2^1$ | Axial fiber family material parameters | 0.36 kPa, 13.14 |
| $c_1^2, c_2^2$ | Circumferential fiber family material parameters | 1.24 kPa, 1.83 |
| $c_1^{3,4}, c_2^{3,4}$ | Diagonal fiber families' material parameters | 0.003 kPa, 18.56 |
| $\eta^3, \eta^4$ | Diagonal fiber families' alignment parameter | 26.37<sup>0</sup>, -26.37<sup>0</sup> |
| $\lambda_m, \lambda_0$ | Maximum, minimum contractile stretch | 2.5, 0.2 |

Aggrecan and versican are proteolytically cleaved by ADAMTS1, 4, 5, and 9 (12). *ADAMTS1* and *ADAMTS4* mRNAs had higher levels in the venous wall in microarrays (Fig. 1e, Sup. Table 1) and RNA-ISH demonstrated stronger expression of *ADAMTS1, ADAMTS4, ADAMTS5,* and *ADAMTS9* in the veins. *ADAMTS1* was the most strongly expressed, localizing to venous endothelium and TM, with modest umbilical artery expression seen in the outer TM (Fig. 2a). *ADAMTS9* was similarly expressed as *ADAMTS1*, whereas *ADAMTS4* and *ADAMTS5* mRNAs were restricted to umbilical vein endothelium (Fig. 2a). Neo-epitope antibodies detecting ADAMTS-cleaved aggrecan and versican (anti-NITEGE and anti-DPEAAE respectively) (13–15) showed strong staining throughout the venous TM and in the outer arterial TM, but not the inner arterial TM (Fig. 2b). Thus, proteoglycan accumulation in the inner TM of the umbilical artery results from higher *ACAN* and *VCAN* expression and lower proteolysis. In contrast, lower *ACAN* and *VCAN* expression and greater ADAMTS activity within the umbilical vein may preclude proteoglycan accumulation.

**Figure 2.**
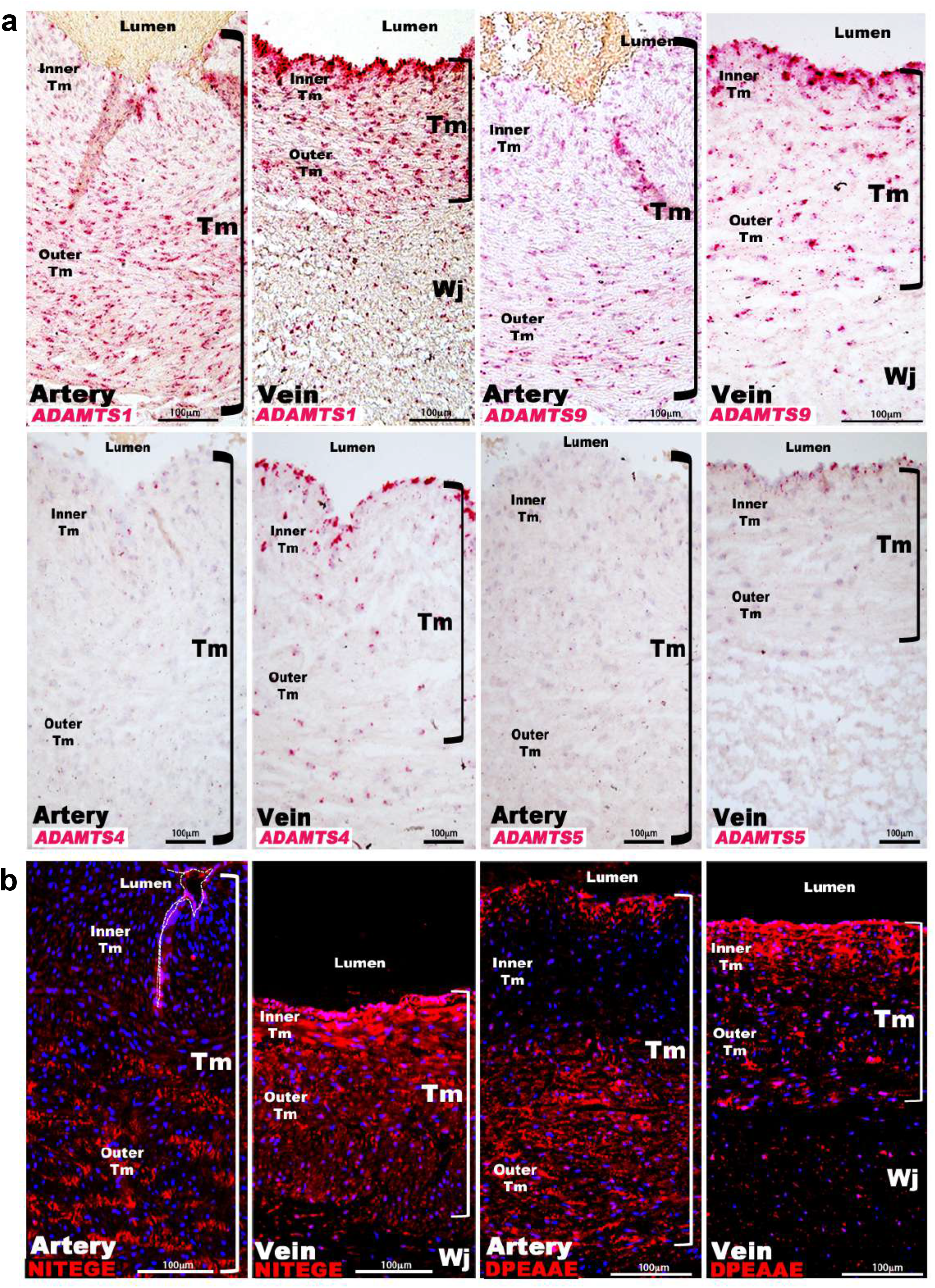
ADAMTS proteoglycanases are selectively expressed and active in the umbilical vein. **(a)** In situ hybridization shows robust *ADAMTS1* and *ADAMTS9* expression in umbilical vein endothelium and TM and in outer arterial TM. *ADAMTS4* and *ADAMTS5* expression were confined to the venous endothelium. **(b)** ADAMTS-cleaved aggrecan (anti-NITEGE, red) and versican (anti-DPEAAE, red) showed ADAMTS activity throughout the venous wall, but only in the outer artery TM. Unlike aggrecan, extensive versican proteolysis is seen in the arterial intima and sub-intima. Wj, Wharton’s jelly. The brackets mark TM boundaries. Scale bars in **a-b**= 100µm.

We postulated that abundant hydrated proteoglycans in the arterial inner TM provided compressive stiffness that could not only prevent kinking and premature occlusion but could potentially facilitate rapid umbilical artery closure at birth. If so, similar adaptations should be present in other mammals. Analysis of umbilical cords from nine large primate and non-primate mammals disclosed similar dimorphism, i.e., umbilical arteries were occluded and had thicker walls with similar infolding of the arterial inner TM (Fig. 3a) and strong alcian blue and CS-staining, contrasting with veins (Fig. 3a,b). Anti-aggrecan and anti-NITEGE stained several animal species, confirming aggrecan abundance in the arterial TM and robust aggrecan cleavage on the venous side (Fig. 3c-d, Sup. Fig. 4a).

**Figure 3.**
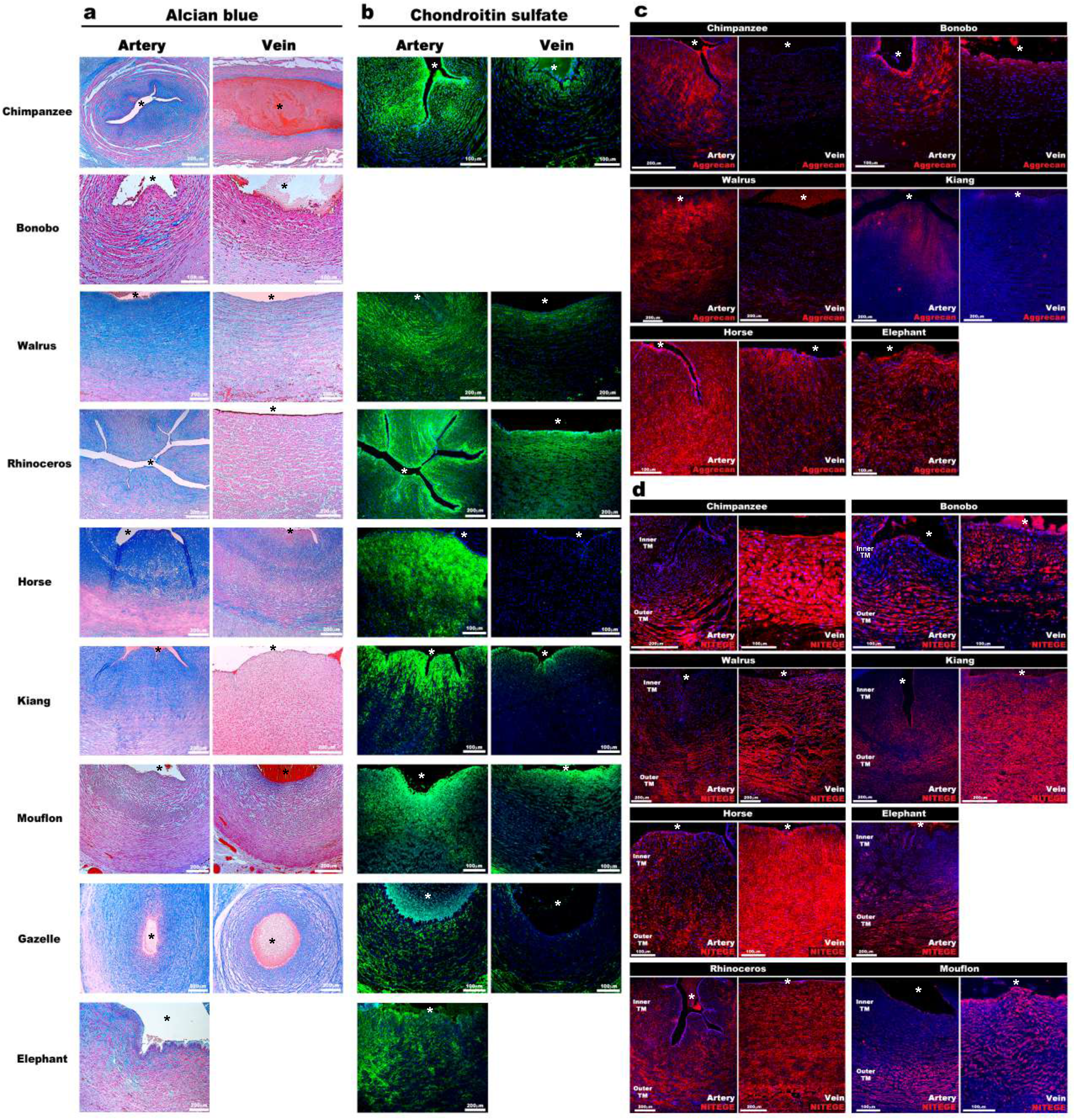
Aggrecan enrichment in the inner umbilical artery tunica media (TM) and its proteolysis in the umbilical vein is a characteristic of large mammals. **(a)** Alcian blue-eosin staining of umbilical cord sections shows proteoglycan enrichment (blue) in the inner arterial TM. The elephant umbilical vein was unavailable. **(b)** Anti-CS immunofluorescence (7D4, green) shows enrichment in the inner arterial TM. Bonobo cords lacked 7D4 reactivity. **(c,d)** Aggrecan and anti-NITEGE immunostaining from reactive species showed aggrecan enrichment in the inner arterial TM and aggrecan proteolysis in the vein and outer artery TM. Scale bars = 100µm and 200µm. * lumen.

Despite uniform staining with SMC markers, co-staining with serine^20^-phosphorylated myosin light chain (pMLC) marking SMCs primed for contraction (16) revealed that umbilical arteries had more contractile SMCs than the vein, predominantly in the outer TM (Fig. 4a, b). Thus, outer umbilical artery SMCs are principally responsive to vasoconstriction stimuli at birth, whereas inner SMCs are relatively non-contractile. We hypothesized that an outer ring of contracting SMCs could drive the CSPG-rich inner TM centripetally, occluding the lumen, and addressed this possibility initially using ex-vivo biomechanical testing of late-gestation mouse umbilical vessels (Fig. 4c, Sup Fig. 5). Umbilical arteries had a smaller lumen, as expected (Sup Fig. 5b) and exhibited lower distensibility and especially extensibility under passive conditions compared to the umbilical veins. Umbilical arteries constricted significantly (30-50% reduction in measured outer diameter at 25 mmHg fixed pressure), causing complete luminal occlusion verified by optical coherence (OCT) imaging, which was not observed in the veins (Fig. 4c). Cross-sectional area measurements at fixed lengths revealed wall volume reductions during vasoconstriction (Sup. Fig. 5a), less in the umbilical vein (~35%) than the umbilical artery (~50%), suggesting fluid exudation from the wall under forceful SMC contraction, especially from the GAG-rich inner layer.

**Figure 4.**
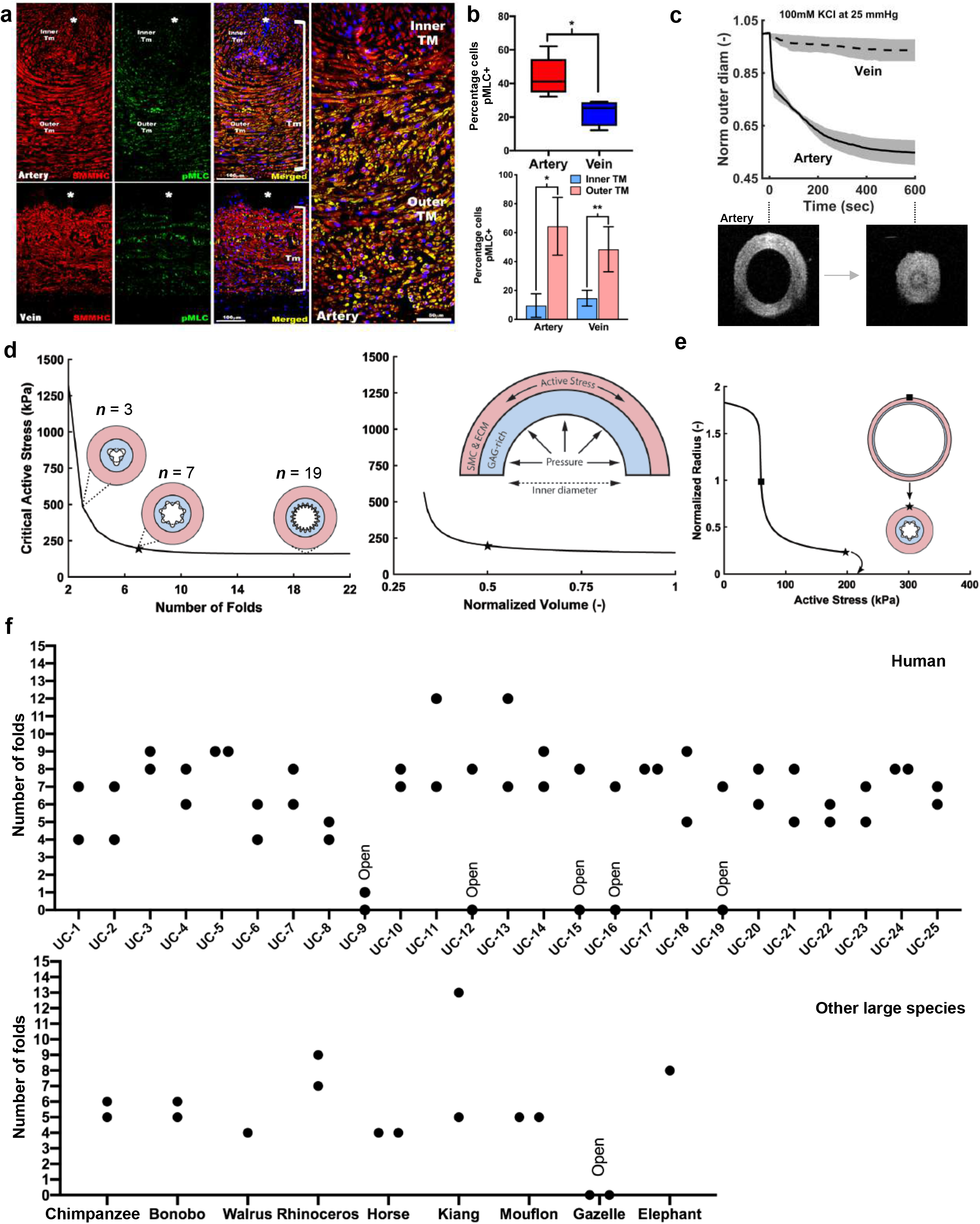
Contraction-induced buckling ensures effective closure of the umbilical artery at birth. (a) Smooth muscle myosin heavy chain (SMMHC-red) and serine-20 phosphorylated myosin light-chain (pMLC-green) show more contraction-primed SMCs in the outer arterial TM (white brackets) than the umbilical vein. Scale bars are 100 µm. (b) Quantitation of pMLC^+^ SMCs in the artery (red) and vein (blue) (top, n=5 arteries, 4 veins, whiskers indicate minimum and maximum values, *, p<0.05), inner and outer TM of both reveal similar distributions but greater activity in the artery (bottom, n=3 arteries, 4 veins, error bars indicate mean ± S.D. *, p<0.05; **, p<0.01). (c) Differential contraction of murine umbilical artery and vein stimulated by 100 mM potassium chloride (KCl) under biaxial loading confirms greater contractility in the artery, with OCT images prior to and following contraction-induced arterial closure. **(d)** Computational simulations of a bilayered artery with contractile SMCs in the outer layer and swollen inner layer: critical contractile stress values leading to buckling for (*left*) different numbers of folds for a normalized inner layer volume of 0.5 and (*right)* decreasing values of normalized volume of the inner layer for 7 folds. **(e)** Normalized inner radius as a function of contractile stress for inner layer volume change of 0.5 and 7 folds. The states for the inflection (square) and critical active stress (star) are illustrated by the schematics; complete closure achieved with contraction-induced buckling. All simulations were run for 25mmHg pressure. Due to the linear stability analysis, the amplitude of the folds in the buckled schematics is illustrative. **(f)** Number of buckles observed in human (top, indicated as UC1 −25) and other large mammalian (bottom) umbilical arteries. Both arteries per cord were included. Open vessel lumens are indicated where observed.

These biomechanical tests informed a novel computational model of the umbilical artery incorporating its complex bilayered, multi-constituent structure (GAG-rich inner layer and SMC- and collagen-rich outer layer; Fig. 4d) and multiaxial mechanical loading: specifically, axial extension, luminal pressurization, active contraction by SMCs, and intramural swelling of the inner layer that regulates tissue volume locally based on GAG content. Nonlinear regression of biaxial mechanical data from passive tests of murine vessels identified best-fit values of the material parameters in the constitutive model for the umbilical artery (Sup. Table 3) while data from active contraction studies guided the selection of the associated active constitutive parameters (Sup. Table 3). Model-based parametric studies examined combinations of different levels of GAG-driven swelling and SMC-generated active stress to identify their roles in umbilical artery closure at different levels of fixed luminal pressure. Increasing inner layer swelling in the absence of active outer layer stress narrowed the lumen at a fixed pressure, as expected given the constraining effect of the outer stiff passive matrix (Sup. Fig. 6a). This trend reversed in the presence of active stresses, with increasing inner layer GAGs able to oppose vasoconstriction if overall wall volume remained constant (Sup. Fig. 6b). Thus, increased inner layer swelling attenuates the ability of SMC contraction to prematurely reduce luminal radius, as revealed by varying the active stress parametrically for different fixed values of inner layer swelling. Importantly, the model predicted a sharp transition from a widely patent to a narrowed lumen due to small changes in active stress at lower values of swelling whereas radius changes were more gradual at higher values of swelling (Sup. Fig. 6b). This transition, at which a decrease in volume of the inner layer associates with larger or smaller luminal radii for values of active stress below or above T_*act*_ ≅ 60 kPa (a key parameter of active stress generation) appears to be close to the *in vivo* value. Hence, consistent with *ex vivo* findings (Sup. Fig. 5), it appears that inner layer volume loss during increased SMC contraction aids vessel narrowing. Regardless, the inner radius reached nearly constant values for increasing levels of active stress (Sup. Fig. 6b). Thus, contraction alone is insufficient to occlude the vessel.

Given the consistent histological finding of inner TM infolding following birth, we modeled the biomechanics of superimposed inner layer buckling in the bilayered arterial model. Buckling can release energy stored in the inner layer during vasoconstriction-induced compression, thereby reducing the structural stiffness resistance to SMC contraction. This analysis parametrically considered possible perturbations to cylindrical geometry achieved at various levels of fixed luminal pressure and different values of swelling and actively generated wall stress. Examining the influence of the number of inner layer folds for different values of swelling disclosed higher inward buckling probability with more folds (Fig. 4d). Since T_*act*_ needed to cause buckling tended to plateau at 7 folds, we used 7 folds subsequently for illustrative purposes. T_*act*_ needed to cause buckling decreased for a more swollen inner layer (Fig. 4d, Sup. Fig. 6c) and increased exponentially with inner layer volume loss. This finding was likely due to the less negative values of circumferential wall stress in the inner layer occurring with shrinkage (Sup. Fig. 6d). We found that an arterial wall consisting solely of SMCs and uniform matrix maintained a mean positive circumferential stress in the inner layer during contraction, preventing buckling. Thus, a delicate biomechanical balance – reduced inner layer volume allows a smaller radius to be achieved via SMC contraction prior to buckling (Sup. Fig. 6b), thus aiding closure, yet excess volume reduction of the inner layer increases the active stress requirement for buckling and achieving complete vessel closure (Fig. 4d). The umbilical artery can reduce its cylindrical radius dramatically at T_*act*_ near a basal value of ~60 kPa, progressing to buckle and close via a subsequent near-maximal contraction (Fig. 4e). In agreement with the computational modeling, 20/25 of human umbilical arteries had 4 or more buckles whereas those with fewer than 4 or no buckles were patent (Fig. 4f). Other large mammalian species showed a similar phenomenon (Fig. 4f).

Mouse umbilical cords also demonstrated vascular dimorphism (Figure 5a), suggesting that genetically modified mice could provide mechanistic insights into proteoglycan dynamics and its impact. Aggrecan and versican immunofluorescence showed strong staining in the mouse umbilical artery inner TM and adventitia, with weaker staining in the veins (Fig. 5b). *Acan, Vcan* and *Adamts1,4,5,9* RNA-ISH at early (12.5) and late (E18.5) gestational stages showed that *Acan* and *Vcan* were strongly expressed in the umbilical arteries (Supplemental Fig. 7a). *Adamts1* was the most highly expressed proteoglycanase in the mouse umbilical vein pre-parturition (E18.5) (Supplemental Fig. 7a), evidenced by strong β-gal staining in endothelium and TM of *Adamts1*^lacZ/+^ embryos had staining that was absent in the arterial TM (Fig. 5c). Although *Adamts9* mutant embryos were observed to have short umbilical cords, abnormal umbilical artery development, and to die by 14.5 days of gestation (E14.5) (17), umbilical cord development was not previously investigated in mutants of the two genes implicated here as potentially critical for umbilical cord vascular dimorphism, *Acan* and *Adamts1*.

**Figure 5.**
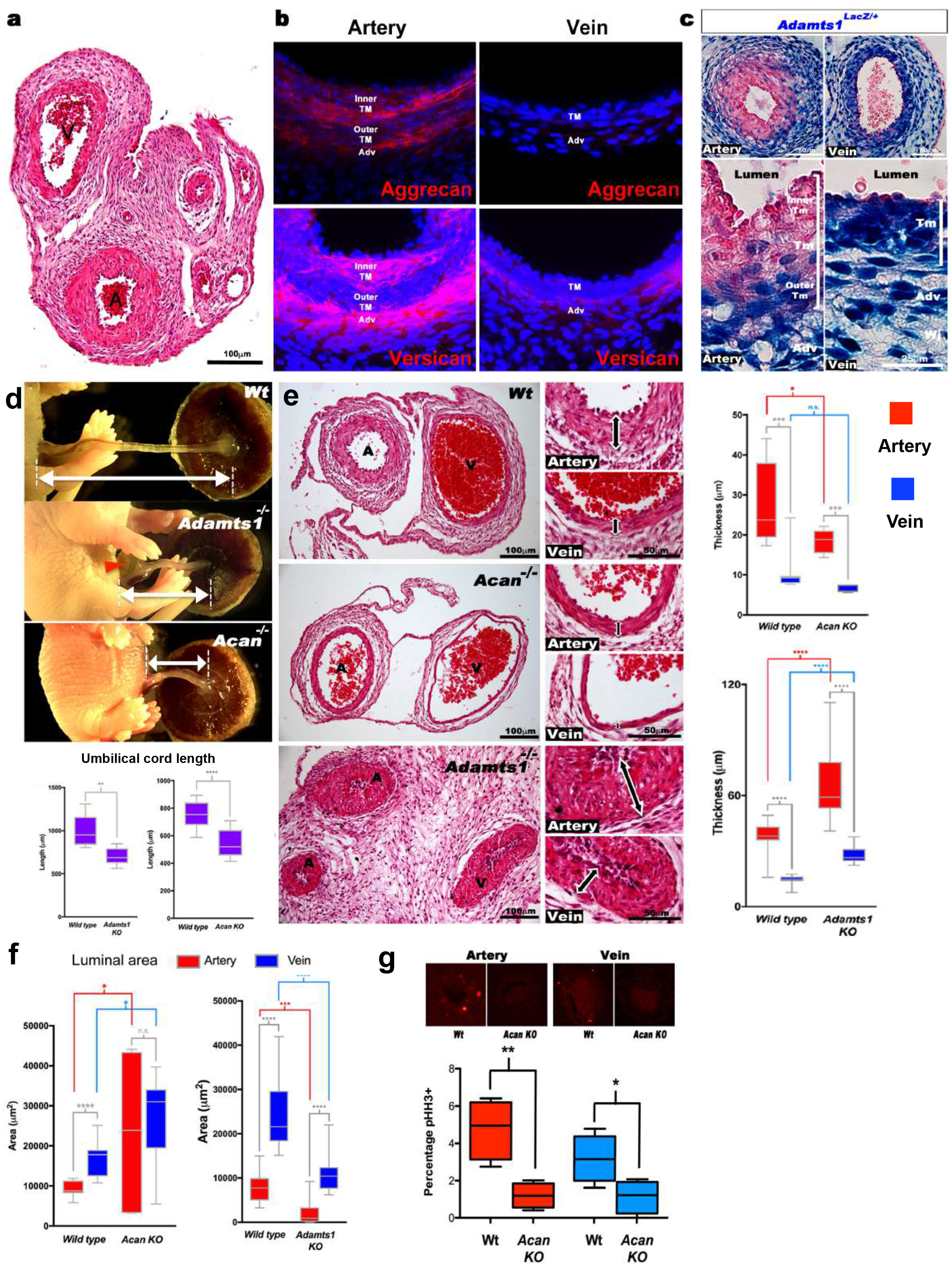
Defective morphogenesis in *Acan* and *Adamts1* null mouse umbilical cords. **(a)** H&E staining of E18.5 wild-type cords showing thicker umbilical arterial (A) and thinner venous (V) wall. **(b)** Aggrecan and versican localization (red, DAPI counterstain blue) in E18.5 wild-type cords showing staining in the arterial inner TM and adventitia but not the vein. **(c)**β-gal (blue) and eosin (red) staining of E18.5 *Adamts1^LacZ^*^/+^ cord showing strong *Adamts1* expression in venous endothelium and TM and outer artery TM. **(d)** Shorter umbilical cords in E18.5 *Adamts1*^*−/−*^ and *Acan^cmd/cmd^* embryos compared to wild type. Red arrowhead indicates an omphalocele in *Adamts1^−/−^* embryos (n=5 umbilical cords each, whiskers indicate minimum and maximum values, **, p<0.01; ****, p<0.0001 **(e)** H&E staining of E18.5 wild type, *Acan^cmd/cmd^* and *Adamts1*^*−/−*^ cord cross-sections showing thinner walls in *Acan^cmd/cmd^* and thicker walls in *Adamts1^−/−^* vessels (n=7-11 umbilical cords each, whiskers indicate minimum and maximum values, *, p<0.05; **, p<0.01; ***, p<0.001; ****, p<0.0001) **(f)** Luminal area quantification of E18.5 Acan^cmd/cmd^ and Adamts1^−/−^ cord vessels showing larger lumens in Acan^cmd/cmd^ umbilical cords and smaller lumens in Adamts1^−/−^ (n=7-11 umbilical cords each, whiskers indicate minimum and maximum values, *, p<0.05; **, p<0.01, ***, p<0.001, ****, p<0.0001). (**g**) Phospho histone-H3 (pHH3) staining shows significantly fewer proliferating cells in Acan^cmd/cmd^ umbilical vessels (n=4 cords each, whiskers indicate minimum and maximum values, **, p<0.001; *, p<0.05). Scale bars = 100µm in **a**, 25µm in **c**, 100µm and 50µm in **e**.

Neither *Acan^cmd/cmd^* (10, 18) nor *Adamts1^−/−^* embryos survived past birth. *Acan* mutants are thought to succumb to respiratory failure resulting from soft tracheal cartilages and ribs, whereas the cause of *Adamts1^−/−^* lethality is unknown. Neither mutant showed intrauterine growth retardation, and intrauterine death was infrequent, suggesting adequate cord circulation. At E18.5, *Acan^cmd/cmd^* and *Adamts1^−/−^* mutants each had significantly short umbilical cords (Fig. 5d) demonstrating their requirement for proper umbilical cord development. Histology showed thinner vascular walls in *Acan^cmd/cmd^* cord vessels, and conversely, thicker vascular walls in *Adamts1^−/−^* embryos (Figure 5e). At earlier developmental stages (e.g., E14.5), lack of aggrecan neither impaired umbilical cord growth (Supplemental Fig. 7b), nor circumferential SMC orientation (Supplemental Fig. 7c), which occurs around E13.5 and is defective in *Adamts9* mutants (17). Thus, *Acan* and *Adamts1* are involved in cord vessel development from early gestation, but their functions manifest near parturition. The arterial lumen was not reduced in *Acan^cmd/cmd^* mice relative to wild-type whereas arterial and venous lumens were smaller in *Adamts1^−/−^* mice, indicating that their thicker vascular walls compromised luminal diameter.

Phospho-histone H3 staining revealed fewer proliferating cells in *Acan ^cmd/cmd^* umbilical cords at E18.5 (Fig. 5j). Immunostaining for SMC markers smooth muscle α-actin (SMA), smooth muscle myosin heavy chain (SMMHC), and phosphorylated myosin light chain (pMLC) showed weaker intensity in *Acan^cmd/cmd^* umbilical arteries compared to wild type (Fig. 6a, b). In contrast, *Adamts1^−/−^* umbilical vessels showed stronger SMA, SMMHC and pMLC staining than wild-type littermates and overgrowth of the arterial and venous walls (Fig. 6c). Intriguingly, endomucin, a specific venous endothelium marker (19), was located in *Adamts1^−/−^* umbilical arterial endothelium (Fig. 6c). Immunostaining of E17.5 *Adamts1^−/−^* umbilical cords indicated a crucial role for ADAMTS1 in regulating proteoglycan dynamics. We observed robust accumulation of aggrecan and versican in the mutant vein and in the outer TM of the mutant artery (Fig. 7a-d) and severe loss of the aggrecan and versican neo-epitope staining (Fig. 7a-d).

**Figure 6.**
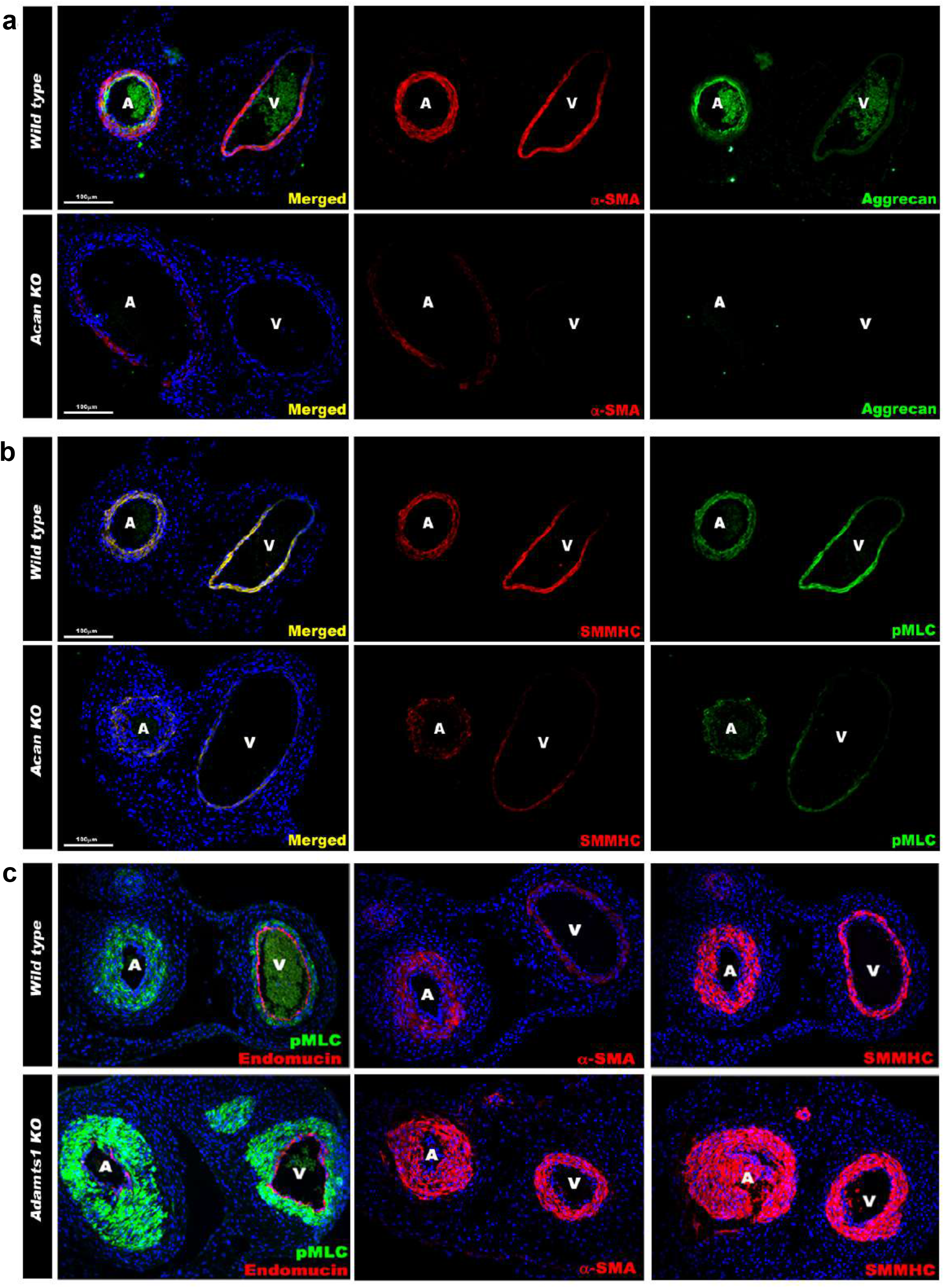
Contrasting smooth muscle cell (SMC) phenotype modulation in *Acan* and *Adamts1*-deficient umbilical vessels. **(a**) Aggrecan (green) and a-SMA staining (red) in E18.5 umbilical cords shows loss of aggrecan and α-SMA staining in the *Acan^cmd/cmd^* vessels. **(b**) Smooth muscle myosin heavy chain (SMMHC, red) and phosphorylated myosin light chain (pMLC, green) staining in E18.5 umbilical cords showing loss of contractile SMCs in the *Acan^cmd/cmd^* vessels. **(c**) pMLC (green), endomucin (red), α-SMA (red, center panels) and SMMHC (red, right-hand panels) stainings shows blunted dimorphism of the umbilical vessels with stronger expression of differentiated SMC markers in *Adamts1*^−/−^ umbilical cords and acquisition of endomucin, a venous endothelium marker, by arterial endothelium. Scale bars= 100µm in **a-c**.

**Figure 7.**
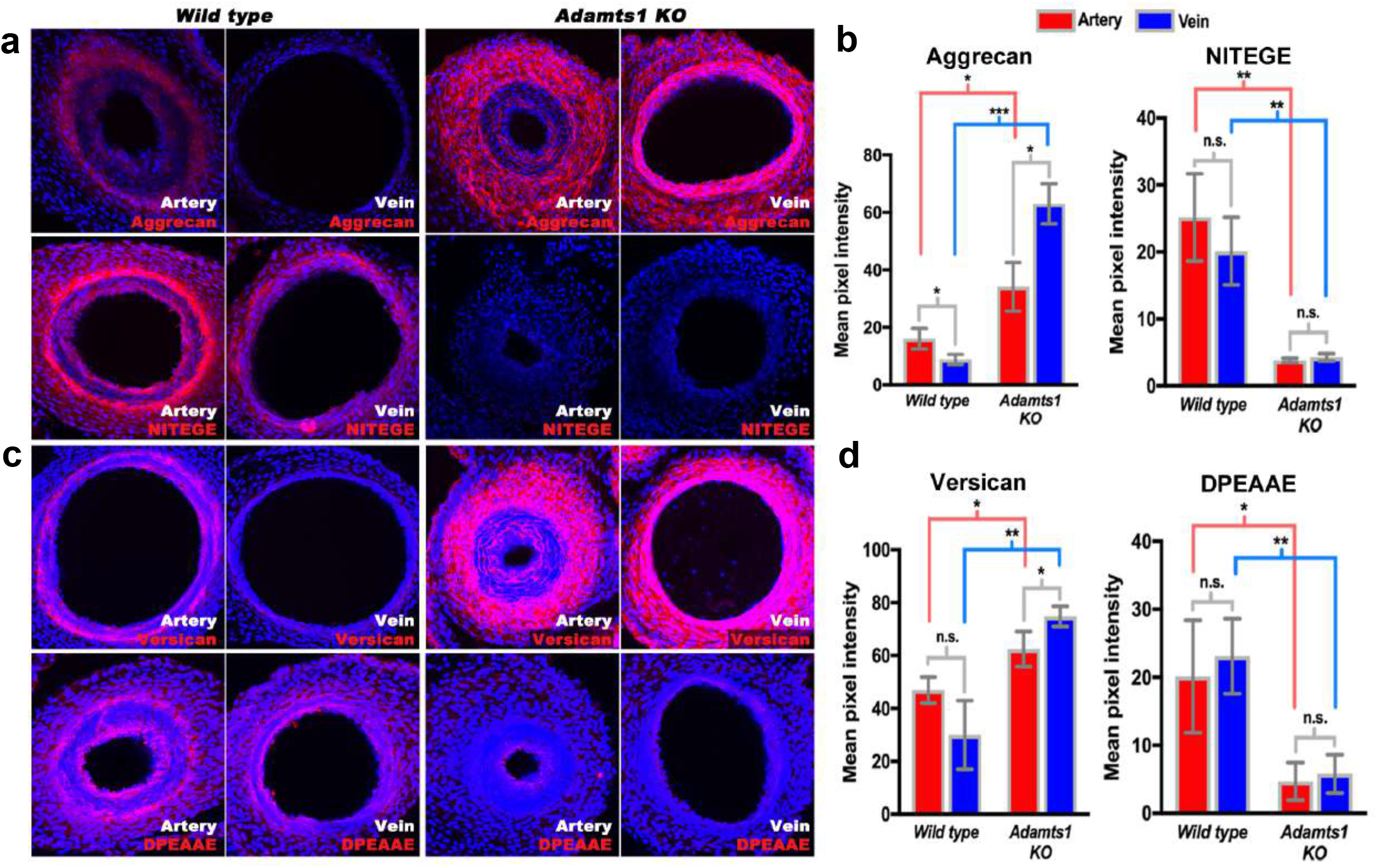
Reduced aggrecan and versican proteolysis in *Adamts1*−/− umbilical cords. **(a,b)** E17.5 *Adamts1*−/− umbilical cords show increased aggrecan and reduced anti-NITEGE staining in **(a)**, quantified in **(b)**(n=3 cords each genotype, error bars indicate mean ± S.D.*, p<0.05; **, p<0.01; ***, p<0.001. **(c,d)***Adamts1*−/− umbilical cords show increased versican **(c)** and reduced anti-DPEAAE staining) quantified in **(d)**(n=3 cords each genotype, error bars indicate mean ± S.D. *, p<0.05; **, p<0.01.

## DISCUSSION

We report two distinct umbilical cord vascular specializations that may facilitate rapid umbilical artery occlusion at birth; a distinct proteoglycan-rich inner arterial TM, generating a bilaminar arterial wall, and selective contraction of smooth SMCs in the outer lamina. The rounded SMC of the arterial inner TM may be specialized for CSPG production rather than contraction, consistent with Sox9 expression, a function they exert prior to delivery. During delivery, lack of pMLC staining suggests that despite SMC marker expression, they play a passive role. Biomechanical testing and computational analysis confirm that selective proteoglycan enrichment in the inner TM ensures that contracting SMCs in the outer TM can effectively occlude the arterial lumen at birth. Histologic and computational analysis ed buckling of the inner TM and fluid redirection into the resulting TM protrusions as critical mechanisms resulting from specialization of the inner and outer arterial TM. By *in silico* simulations of human umbilical arteries with modulation of the contractile outer layer and proteoglycan-rich inner core, we demonstrate that complete occlusion can be achieved.

Our work suggests that a principal mechanism governing umbilical cord vascular dimorphism resides in extracellular matrix, namely, differentially regulated dynamics of aggrecan and versican, which then modulate SMC differentiation. (Fig. 8a-c). Other mammalian umbilical vessels examined, from animals as large as the walrus and elephant to as small as mice showed similar CSPG and aggrecan modulation as humans. Although *Vcan* mutant mice die before allantois development is completed and could not be studied, *Acan* and *Adamts1* mutants demonstrated their mechanistic contribution to the observed dimorphism. Our emphasis on aggrecan in the inner layer, with its abundant GAGs and their high fixed charge density, was relevant to the computational model’s findings regarding the importance of inner layer swelling and buckling in response to SMC contraction.

**Figure 8.**
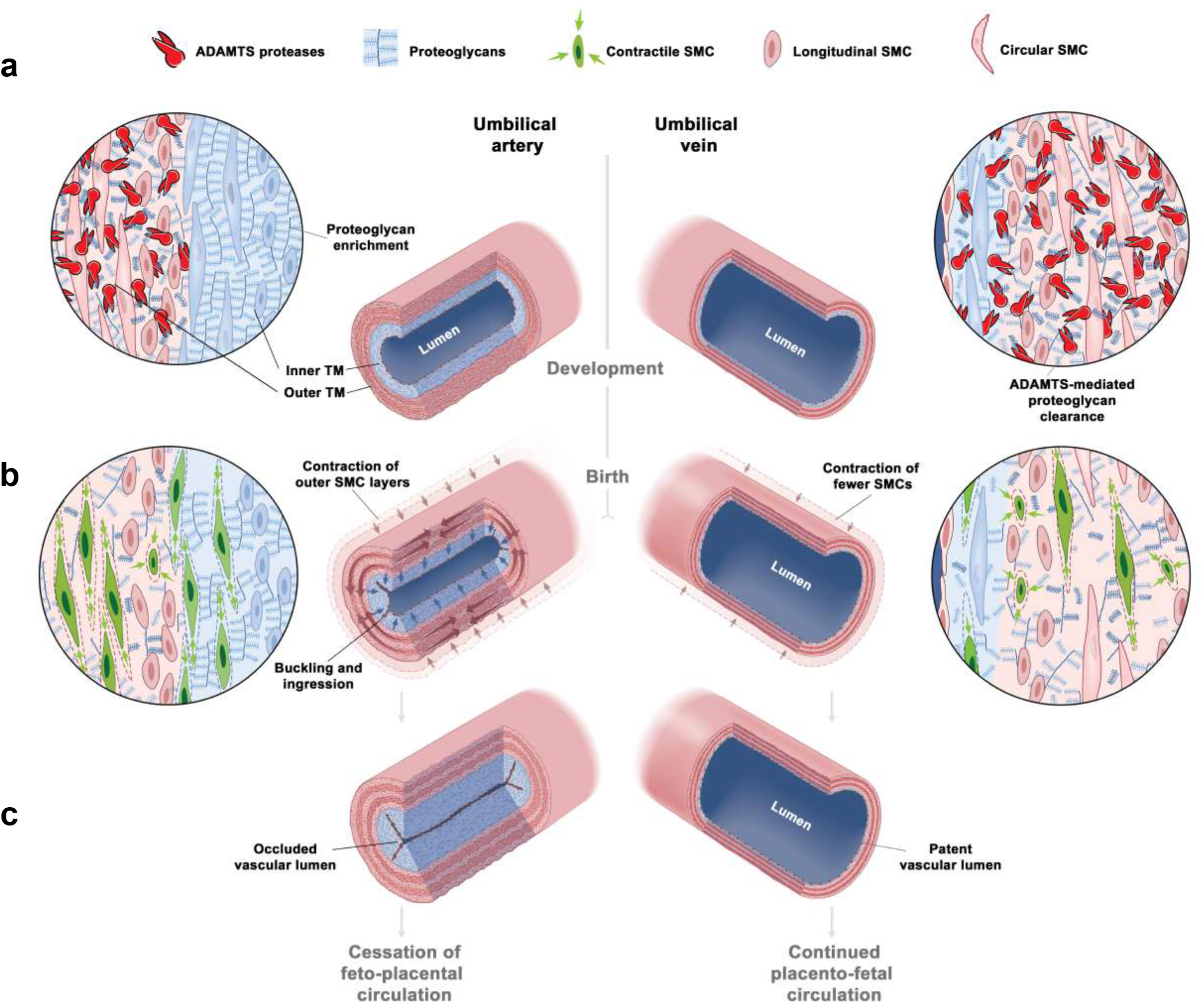
The unique bilayered design of the umbilical artery underlie its rapid occlusion at birth. **(a)** Differential expression of ADAMTS proteases and large CS-proteoglycans results in a bilayered artery with a hydrated proteoglycan-rich inner layer and most contractile SMC in the outer layer, contrasting with the umbilical vein. **(b)** At birth, SMC contraction in the outer layer and fluid movement-induced inner layer buckling redirect the inner layer into the lumen. The single layered vein does not undergo buckling. **(c)** Umbilical artery occlusion at birth prevents neonatal exsanguination, whereas the patent vein allows a final transfusion from the placenta.

The umbilical vein wall contains fewer SMCs primed for contraction and a paucity of aggrecan and versican. Hence, SMC activation in the umbilical vein does not occlude the lumen to the same degree as in the arteries, a contention supported by computational analysis. Given the evolutionary pressure to achieve hemostasis in the artery rather than the vein, these findings suggest a highly evolved mechanism for preventing exsanguination of the newborn that is potentially relevant to other embryonic shunts that occlude rapidly at birth.

The computational model was built on a long history of studying murine arteries (20), but was specialized to the umbilical artery’s GAG-rich inner layer and SMC-rich outer layer. Modeling the dynamics of such volume changes would have required a mixture or poro-elastic model and significantly more experimental data, including measurement of layer-specific permeabilities and fixed-charge densities. Instead, we modeled the equilibrated states using a well-accepted approach, wherein the degree of swelling can be adjusted for each simulation (21, 22). Interestingly, prior results by others showed that swelling of an initially unloaded, unilayered cylindrical tube consisting of a neo-Hookean material (which we used to model GAG-rich tissue) increases luminal diameter if the tube is unconstrained (21). The tube must be constrained, such as by a stiff outer layer surrounding the swollen layer if swelling is to decrease luminal diameter (22). The abundance of contractile SMCs in the stiffer outer layer and their basal tone may in fact enhance outer layer stiffness and hence buckling appears to be essential to augment contraction-induced closure of the umbilical artery (cf. (23)).

The observed architecture of the umbilical cord is likely to have supported survival of mammalian species, humans included, thousands of years before formal obstetric involvement. Cord clamping, a relatively modern innovation, supplements a natural and apparently conserved physiologic process in the transition from fetal to neonatal life. It can therefore be seen as an adjunct offering control over fetal well-being and an option to resuscitate quickly if needed, rather than a strict necessity. Its contemporary use has the sanction of convention and it offers benefits in reducing maternal and fetal mortality and morbidity, particularly where immediate neonatal resuscitation is required or severe maternal hemorrhage occurs. In contrast, evolution has devised a mechanism that works nearly all the time, which suffices, since evolutionary success is not about minimizing poor outcomes, but ensuring survival of a significant majority.

## Methods

### Human and large mammal cords

25 human umbilical cords were obtained from uncomplicated term pregnancies either after vaginal birth (N=13), or Cesarean section for obstetric indications (N=12), i.e. malpresentation or repeat Cesarean section. The samples were collected under an IRB exemption (exemption 4) from Cleveland Clinic for use of discarded tissue without patient identifiers. These cords were used for histological/immunohistologic analysis, *in situ* hybridization, and transcriptomics of inner *vs* outer umbilical artery TM. For microarray analysis of umbilical cord artery versus vein, human umbilical cords were collected separately through the National Children’s Study (NCS). Animal cord sections were provided by Disease Investigations, Institute for Conservation Research, San Diego Zoo Global from the Benirschke archive. Additional details including antibodies, RNA probes and microarray analysis are provided in the Online Supplement.

### Mutant mice

The *Adamts1* transgenic allele (*Adamts1*^tm1Dgen^) referred to herein as *Adamts1*^-^, was produced by insertion of an IRES-lacZ cassette into intron 1 of *Adamts1* using homologous recombination in mouse embryonic stem cells (24). These mice were rederived by the Case Transgenic Core from frozen embryos purchased from the European Mutant Mouse Archive (J:82809). The *Acan*^cmd-Bc^ allele was previously described (18). The line was back-crossed to C57BL/6 (10) and referred to herein as *Acan*^cmd/cmd^. Mice were handled under standard conditions under approved IACUC protocols at the Cleveland Clinic (IACUC protocol nos. 18-1996 and 18-2045) and University of Chicago (IACUC protocol no. 43751).

### Biomechanical and computational analysis

E18.5 umbilical artery and vein were obtained at Yale University (IACUC protocol no 2018-11508) and mounted within a custom computer-controlled biaxial device for biomechanical testing (25). Vessel maintenance, pre-conditioning, biaxial protocols and data collection are described in the Online Supplement. The umbilical artery was modeled computationally as a thick-walled, bi-layered cylindrical tube subjected to swelling of the GAG-rich inner layer and active contraction of the smooth muscle-rich outer layer; the model also phenomenologically included a passive contribution of matrix as revealed by biomechanical tests. Additional details are provided in the Online Supplement.

## Supporting information

Online microarray data supplement-inner versus outer umbilical artery

Online microarray data supplement- umbilical artery versus vein

## Acknowledgements

We acknowledge the Paul Scherrer Institut, Villigen, Switzerland for provision of synchrotron radiation beamtime at the TOMCAT beamline X02DA of the SLS and would like to thank Goran Lovric for assistance. We are grateful to the Disease Investigations team at San Diego Zoo Global for use of sections from the Benirschke archive, and to the late Dr. Kurt Benirschke for collecting the animal umbilical cords used in this study.

