## Supplemental Methods, Figures and Table for "Vascular dimorphism ensured by regulated proteoglycan dynamics favors rapid umbilical artery closure at birth"

**Online supplement**, including

Supplemental methods
Supplemental figures

**Supplemental data online**

Supplemental array data set 1: Microarray data Artery vs Vein
Supplemental array data set 2: Microarray data Artery Inner TM vs outer TM
Movie S1: Synchrotron-based phase contrast micro-CT of human umbilical artery
Movie S2: Synchrotron-based phase contrast micro-CT of human umbilical vein

#### SUPPLEMENTAL METHODS

##### Human tissue

After delivery of the neonate, the umbilical cord was doubly clamped and cut. The time from delivery to cord clamping varied between deliveries and in general ranged from 5-30 seconds. Two additional clamps were placed on the cord after a fetal blood sample was obtained for clinical typing, and scissors were used to cut between these two clamps to obtain a segment of umbilical cord for analysis. Umbilical cord samples were fixed in 4% paraformaldehyde for 48 h and embedded in paraffin. For microarray analysis of umbilical cord artery versus vein, human umbilical cords from separate individual patients were collected as a part of the National Children's Study (NCS) (UHCMC IRB# 01-11-28). Immediately upon neonatal delivery, cords were sectioned into 1-inch segments (for a total of 3 segments per cord at 0 h) and flash frozen with liquid nitrogen. *RNA Later-ICE* (Ambion) was used to maintain nucleic acid viability during any freeze-thaw of tissue. Veins and arteries of the umbilical cords were dissected from each segment for discrete homogenization. RNA from homogenized vein and artery tissue was extracted using Qiagen's RNeasy nucleic acid isolation kit. For microarray analysis of the inner versus outer arterial tunica media, human umbilical cords were dissected immediately upon neonatal delivery and sectioned into 1-inch segments (for a total of 3 segments per artery) and the inner and outer tunica media were carefully dissected under a dissecting microscope and flash frozen in liquid nitrogen in Trizol reagent (Ambion). RNA from homogenized artery tissue was extracted using the chloroform-isopropanol precipitation method.

##### Histological and immunostaining

Seven (7) μm-thick paraffin sections were collected using a Leica RM 2255 microtome and stained with hematoxylin and eosin, Alcian blue, or Masson's trichrome after deparaffinization.

For immunostaining, sections were deparaffinized, and boiled in citrate buffer (10mM citric acid, 0.05% Tween 20, pH 6.0) for 90 seconds for antigen retrieval, washed with PBST and blocked in 10% normal goat serum before staining with the following primary antibodies and incubated overnight at 4°C: Cy3-conjugated smooth muscle $\alpha$-actin ($\alpha$-SMA) (1:400, Sigma C6198), anti-smooth muscle myosin heavy chain (SMMHC) (1:400 Kamiya Biomedical, MC352), anti-serine$^{20}$-phosphorylated myosin light chain (pMLC) (1:200, Millipore, 06-570), anti-chondroitin sulfate (7D4) antibody (Sorrell JM et al., J Histo. Chem 1990 (Ref# 10), a generous gift from Drs. Bruce Caterson and Clare Hughes, Cardiff University, (1:200), FITC-conjugated anti-heparan sulfate (10E4) antibody (1:200, US biological H-1890), polyclonal anti-versican (anti-VC; 1:400). Alexa-488 or Alexa-568 conjugated secondary antibodies against rabbit and mouse IgG were used at 1:400 dilution. Vectashield (H-1200) mounting medium containing DAPI was used for staining nuclei. All images were taken using an Olympus BX51 microscope connected to a Leica DFC 7000T camera under bright field or fluorescent microscopy modes. Multi-channel fluorescent images were merged using the NIH Image J software.

**Synchrotron-based phase contrast micro-CT**

Imaging of arteries and veins from three different umbilical cords was performed at the X02DA TOMCAT beamline of the Swiss Light Source (SLS) at the Paul Scherrer Institute (Villigen, Switzerland). A 4x magnifying objective was used, resulting in a field-of-view of 4.2 x 3.5 $mm^2$ and an effective pixel size of 1.63 x 1.63 $\mu m^2$. For each scanned vessel, a stack of 1080 tomographic images was acquired. Data analysis was performed using NIH Image J and Amira®. Amira® allowed for visualization of the vessels from any angle and the images in Figure 1 and FigureS1 were created by combining two different tomographic imaging planes at a 90-degree angle. NIH Image J was used for creating the AVI video clips movie S1 and S2.

**Microarray analysis of human umbilical cords**

NCS samples to evaluate changes in expression in artery versus vein were run on the Affymetrix Hu-Gene U219 microarray Peg-plate; 150 ng of input RNA from each sample was labeled using an Affymetrix 3' IVT labeling protocol. Samples hybridized to the PEG arrays were washed, stained, and scanned by the Affymetrix Gene Titan. RNA from 2 of the 4 cords was also evaluated on the Affymetrix Hu-Gene 1.1 ST Peg-plate microarray, using the Whole Transcriptome (WT) labeling protocol also starting with a 150 ng of input total RNA. The hybridized PEG Arrays were washed, stained, and scanned by the Affymetrix Gene Titan according to standard protocols. RNA (150 ng) from Inner and outer TM samples was labeled using Affymetrix's WT PLUS protocol. Labeled samples were hybridized overnight to Affymetrix Hu-Gene 2.0 ST microarray cartridges. The sample was removed, then the microarray cartridges were washed and stained on the Affymetrix GeneChip Fluidics Station450 and scanned by the Affymetrix GeneChip Scanner 3000. Total RNA was labeled using Affymetrix's FLASH Tag Protocol. Labeled samples were hybridized overnight to Affymetrix GeneChip miRNA 2.0 Array, which interrogates all the mature human miRNA sequences in miRBase Release 20microRNAs. The sample was then removed and the microarray cartridges were washed and stained on the Affymetrix GeneChip Fluidics Station450 and scanned by the Affymetrix GeneChip Scanner 3000. Gene expression analysis for each microarray study was performed using Affymetrix's Transcriptome Analysis Console (TAC 4.0) through the RMA-SST sketch algorithm and R version 3.5.2. Fold changes were calculated by an empirical Bayes ANOVA method through the TAC 4.0 software. Parameters for gene expression changes include a p-value ≤0.05 and a fold change value ≥ 1.5 and ≤ -1.5. Data collected from samples labeled by different labeling protocols, obtained using different scanners or hybridized to different arrays were analyzed as independent data sets.

**RNA in situ hybridization**

Six (6) μm paraffin sections were probed with RNAscope probes according to manufacturer's guidelines. The following probes were used to detect: *ACAN* (ACD Cat. No. 506841), *VCAN-E8* (ACD Cat. No. 452241), *ADAMTS1* (ACD Cat. No. 524501), *ADAMTS4* (ACD Cat. No. 537341), *ADAMTS5* (ACD Cat. No. 427611), *ADAMTS9* (ACD Cat. No. 445321), *Acan* (ACD Cat. No. 439101), *Vcan-E8* (ACD Cat. No. 428321), *Adamts1* (ACD Cat. No. 463361), *Adamts4* (ACD Cat. No. 497161), *Adamts5* (ACD Cat. No. 427621), *Adamts9* (ACD Cat. No. 400441). The probes were detected using RNAscope 2.5 HD Red reagent kits (ACD Cat. No. 322350).

**Biomechanical testing of mouse umbilical arteries and veins**

The umbilical cord was excised and separated from the placenta from wild-type C57BL/6J mice at embryonic day E18.5. The umbilical artery and vein were identified by opening the abdomen of the embryo and locating the artery connected to the iliac artery and vein connected to the inferior vena cava through the ductus venosus. After separating and cleaning the umbilical arteries and veins from any excessive adipose tissue, the intact vessels were cannulated on custom-drawn glass micropipettes, secured with silk sutures at each end, and mounted within the custom computer-controlled biaxial test device. The vessels were submerged in Krebs-Ringer bicarbonate solution (Krebs) and oxygenated with 95% $O_2$ and 5% $CO_2$ while maintained at 37$^o$C. The umbilical artery lumen closed immediately after separating the umbilical cord from the placenta and was relaxed by waiting 5-15 minutes at 10 mmHg and by washing out with Krebs up to 3 times. The vessels were then subjected to two isobaric (luminal pressure of 5 mmHg, then 10 mmHg) - axially isometric (fixed in vivo axial stretch) contractions by adding 100 mM KCl to ensure viability of the SMC. The transmural organization of the vessel wall was monitored during different stages of the contraction using an optical coherence tomography (OCT) system having an axial (depth) resolution <7 microns and lateral resolution of 8 microns (Callisto Model, Thorlabs, Newton, NJ). For the subsequent passive tests, the normal Krebs was replaced with a $Ca^{2+}$-free Krebs to ensure passive behavior. The vessels were then preconditioned via four cycles of pressurization (artery: 0 - 40 mmHg and vein: 0 – 25 mmHg) while held fixed at their individual in vivo axial stretch. Subsequently, the vessels were subjected to a series of seven biaxial protocols (cyclic pressurization at three different fixed values of axial stretch and cyclic axial stretching at four different fixed values of luminal pressure: artery at 10, 20, 30 or 40 mmHg and vein at 1, 2, 10 or 20 mmHg). Data collected online included outer diameter, luminal pressure, axial length and axial force, which facilitated robust parameter estimation for the constitutive model (Ferruzzi et al., 2015).

**Computational modeling of the umbilical artery**

The traction-free, non-swollen configuration $(R, \Theta, Z)$ is treated as the reference with inner radius, interfacial radius, and outer radius denoted $[A, B, C]$, respectively. Swelling is accounted for via previously outlined methods (Demirkoparan and Pence 2007; Szafron et al., 2017), which leads to a residually stressed, traction-free configuration with corresponding points mapped to $[A^*, B^*, C^*]$ in $(R^*, \Theta^*, Z^*)$. The final loaded configuration, pressurized and axially stretched, is then characterized by $[a, b, c]$ in $(r, \theta, z)$.

Consider a deformation from $(R, \Theta, Z)$ to $(R^*, \Theta^*, Z^*)$ via $\mathbf{F}^* = \mathrm{diag}(\partial R^*/\partial R\,, R^*/R\,, \Lambda_z^*)$, where volume change is imposed by $\det \mathbf{F}^* = v^* = v/V$ with $v^*$ the ratio of volume $v$ in $(R^*, \Theta^*, Z^*)$ to volume $V$ in $(R, \Theta, Z)$. With $v^* = 1$, there is no swelling and the vessel is considered to have no residual stresses; in contrast, $v^* > 1$ (expansion) and $v^* < 1$ (shrinkage) yields self-equilibrating wall stresses in the absence of external loading. Note that $\partial R^*/\partial R = Rv^*/\Lambda_z^* R^*$. Further deformation to the loaded configuration $(r, \theta, z)$ from the swollen configuration $(R^*, \Theta^*, Z^*)$ is then described by $\mathbf{F}^P = \mathrm{diag}(\partial r/\partial R^*\,, r/R^*\,, \Lambda_z)$, where the assumption of incompressibility during transient external loading requires $\det \mathbf{F}^P = 1$, thus $\partial r/\partial R^* = R^*/(\Lambda_z r)$. A multiplicative decomposition of the deformations yields $\mathbf{F} = \mathbf{F}^P\mathbf{F}^* = \mathrm{diag}(Rv^*/\lambda_z r\,, r/R, \lambda_z)$ with $\lambda_z = \Lambda_z^*\Lambda_z$ for convenience. The matrix component $F_{11}$ can also be expressed in terms of the original derivatives, such that $(\partial r/\partial R^*)(\partial R^*/\partial R) = R\lambda_z v^*/r$, which allows us to apply the chain rule and integrate $\int_a^r r\partial r = \int_A^R v^* R/\lambda_z\, \partial R$ to find any radial point $r$ within the vessel wall.

The vessel is assumed to be quasi-equilibrated in any state, such that linear momentum balance requires $\mathrm{div}\,\mathbf{t} = \mathbf{0}$, where $\mathbf{t}$ is the Cauchy stress tensor. Circumferential and axial equilibrium is satisfied identically at each $(r, \theta, z)$. Radial equilibrium requires $\partial r/\partial R + (t_{rr} - t_{\theta\theta})/r = 0$. The Cauchy stress is specialized as $\mathbf{t} = \mathbf{t}^{ex} - p\mathbf{I}$ with $p$ the Lagrange multiplier enforcing incompressibility during transient loading and $\mathbf{I}$ the identity tensor and $\mathbf{t}^{ex}$ the "extra" part of the stress due to deformation and the constitutive response. Integration yields

$$P = \int_a^b (t_{\theta\theta}^{ex} - t_{rr}^{ex})/r dr + \int_b^c (t_{\theta\theta}^{ex} - t_{rr}^{ex})/r dr,$$

where $P$ is the transmural pressure across the vessel wall, with $P > 0$ indicating internal pressurization. We also calculate the overall axial load required for overall equilibrium,

$$L = \pi \int_a^b (2t_{zz}^{ex} - t_{\theta\theta}^{ex} - t_{rr}^{ex}) r dr + \pi \int_b^c (2t_{zz}^{ex} - t_{\theta\theta}^{ex} - t_{rr}^{ex}) r dr + P\pi a^2.$$

See Fig. 4d. The equilibrium problem is solved iteratively for the loaded inner radius $a$ for each luminal pressure $P$ and axial extension $\lambda_z$.

Constitutively, the extra part of the Cauchy stress can be computed from a stored energy density function $W$ for the vessel, with $\mathbf{t} = 2\mathbf{F}(\partial W/\partial \mathbf{C})\mathbf{F}^T/\det \mathbf{F} - p\mathbf{I}$ and $\mathbf{C} = \mathbf{F}^T\mathbf{F}$ the right Cauchy-Green tensor. Due to the microstructure of the umbilical vessels, the GAG-rich inner layer is modeled as a neo-Hookean matrix that can swell (Demirkoparan and Pence, 2007), with

$$W = \frac{\mu_1}{2}\left(\mathrm{tr}(\mathbf{C}) - 3v^{*\frac{2}{3}}\right) \quad \forall\, r < b,$$

with $\mu_1$ a shear modulus for this inner layer. As there is no evidence of collagen with a preferred orientation or smooth muscle cells capable of contraction within the inner layer, it is considered isotropic and passive. Fewer GAGs are present in the outer layer, but we include the possibility of a swellable matrix for illustrative purposes and to provide radial stiffness. The outer layer is then modeled using a modified four-fiber family model for a passive nonlinear stress-stretch behavior and a Rachev-type model for SMC contractility (i.e., active behavior) with a potential function

$$W = \frac{\mu_2}{2}\left(\mathrm{tr}(\mathbf{C}) - 3v^{*\frac{2}{3}}\right) + \sum_{\alpha=1}^{4} \frac{c_1^\alpha}{4c_2^\alpha}\left(\exp\left(c_2^\alpha\left(\lambda^{\alpha 2} - 1\right)^2\right) - 1\right)$$

$$+\mathrm{T}_{act}\left(\lambda_\theta + \frac{1}{3}\frac{(\lambda_m - \lambda_\theta)^3}{(\lambda_m - \lambda_0)^2}\right)\ \forall\, r > b,$$

where $\mu_2$ is shear modulus for the outer layer, $c_1^\alpha$ and $c_2^\alpha$ are material parameters for each fiber family $\alpha = 1,2,3,4$, $\lambda^\alpha = \mathrm{sqrt}(\mathbf{C} : \mathbf{M}^\alpha \otimes \mathbf{M}^\alpha)$, $\mathrm{T}_{act}$ is the magnitude of the active stress, $\lambda_m$ is the stretch at which contraction is maximum, and $\lambda_0$ is the stretch at which contraction ceases (Baek et al., 2007). Each fiber family has an orientation vector $\mathbf{M}^\alpha = \sin(\eta^\alpha)\,\mathbf{e}_\Theta + \cos(\eta^\alpha)\mathbf{e}_Z$ with $\eta^\alpha$ the fiber angle relative to the axial direction. The fiber families are assumed to lie in the circumferential direction ($\alpha = 1, \eta^1 = 90^\circ$), the longitudinal direction ($\alpha = 2, \eta^2 = 0^\circ$), and symmetric diagonal directions about the $z$-axis with $\eta^\alpha$ fit from the experimental data ($\alpha = 3, \eta^3 = \eta^d$ and $\alpha = 4, \eta^4 = -\eta^d$). Parameter values for the passive behavior of the umbilical artery were determined by minimizing the difference between model-generated outputs and data from the biaxial mechanical tests (Table S3) using a nonlinear least squares approach (Ferruzzi et al., 2013).

*Bifurcation Analysis – Basic Approach*

To examine the potential for unstable equilibria (i.e., bifurcations in the solutions) leading to buckling of the wall, we consider an incremental deformation added to the finite deformation (Ogden 1984) with a notation similar to that previously used to describe bifurcation behaviors in growing elastic solids (Li et al., 2011; Moulton and Goriely, 2011), which is mathematically similar to swelling. Quantities related to the intermediate finite deformation are given as $(\cdot)^{(0)}$ while those related to the incremental motions are denoted as $(\cdot)^{(1)}$. The current position $\boldsymbol{x}$ from position $\boldsymbol{x}^{(0)}$ is specified as $\boldsymbol{x} = \boldsymbol{x}^{(0)} + \epsilon\boldsymbol{u}^{(1)}$ with $\epsilon \ll 1$ scaling the displacement $\boldsymbol{u}^{(1)}$, which allows the deformation gradient to be written as $\mathbf{F} = \mathbf{F}^{(0)} + \epsilon\mathbf{H}^{(1)}\mathbf{F}^{(0)}$ with $\mathbf{F}^{(0)} = \mathbf{F}^{P}\mathbf{F}^{*}$ the deformation gradient from the reference to the finitely deformed configuration and $\mathbf{H}^{(1)}$ the incremental displacement gradient with respect to that configuration. The nominal stress $\mathbf{P}$, defined through $\mathbf{t} = \mathbf{FP}$, follows as $\mathbf{P} = \mathbf{P}^{(0)} + \epsilon\mathbf{P}^{(1)}$. As noted previously (Haughton and Ogden, 1978, Sanft et al., 2019), it is convenient to update the reference configuration to the current configuration yielding $\mathbf{P}_0 = \mathbf{P}_0^{(0)} + \epsilon\mathbf{P}_0^{(1)}$, which, along with considering the Lagrange multiplier as $p = p^{(0)} + \epsilon p^{(1)}$, allows us to write $\mathbf{P}_0^{(1)} = \boldsymbol{\mathcal{B}}\colon\mathbf{H}^{(1)} + p^{(0)}\mathbf{H}^{(1)} - p^{(1)}\mathbf{I}$ with $\boldsymbol{\mathcal{B}} = \mathbf{F}^{(0)}(\partial^2 W/\partial\mathbf{F}^{(0)}\partial\mathbf{F}^{(0)})\mathbf{F}^{(0)}$ the fourth-order stiffness tensor calculated in the finitely deformed configuration and $p^{(1)}$ the increment in the Lagrange multiplier. Linear momentum balance then requires $\mathrm{div}\mathbf{P}_0 = \mathbf{0}$, which is satisfied by the finite deformation, leaving $\mathrm{div}\mathbf{P}_0^{(1)} = \mathbf{0}$ to be resolved. We proceed by assuming a form for the incremental displacement describing the buckling as $\boldsymbol{u}^{(1)} = [\mathrm{u}(r,\theta), \mathrm{v}(r,\theta), 0]$, with no incremental motion in the axial direction. Displacements in the $r$ and $\theta$ directions are also not a function of $z$. The incremental displacement gradient then becomes

$$\mathbf{H}^{(1)} = \begin{bmatrix} \mathrm{u}_r & \dfrac{\mathrm{u}_\theta - \mathrm{v}}{r} & 0 \\ \mathrm{v}_r & \dfrac{\mathrm{u} + \mathrm{v}_\theta}{r} & 0 \\ 0 & 0 & 0 \end{bmatrix},$$

where $(\cdot)_r \equiv \partial(\cdot)/\partial r$ and $(\cdot)_\theta \equiv \partial(\cdot)/\partial\theta$. The incremental deformation is assumed to be isochoric, requiring $\mathrm{tr}(\mathbf{H}^{(1)}) = 0$. As there is no axial dependence in the incremental deformation, the equilibrium equations reduce to a system of two differential equations in u, v, and $p^{(1)}$. These functions are assumed to have sinusoidal forms in the buckled configuration, where $\mathrm{u} = f(\mathrm{r})\cos(n\theta)$, $\mathrm{v} = g(\mathrm{r})\sin(n\theta)$, and $p^{(1)} = h(r)\cos(n\theta)$ with $n$ the buckling mode (i.e. the number of folds in the inner portion of the vessel wall). Using the incompressibility condition, it is possible to rewrite $g(r)$ in terms of $f(r)$, and the two equilibrium equations can be combined to eliminate $h(r)$, yielding a single fourth order, ordinary differential equation in $f(r)$ (Haughton and Ogden 1979, Sanft et al., 2019) of the form

$$A_4 f'''' + A_3 f''' + A_2 f'' + A_1 f' + A_0 f = 0,$$

where coefficients $A_0$-$A_4$ are given below and $(\cdot)' = d(\cdot)/dr$. We assume that the incremental tractions on the inner surface are zero with $\mathbf{n} \cdot \mathbf{t}^{(1)} = \mathbf{0}$, which yields two equations

$$\left[B_{1,3} f''' + B_{1,2} f'' + B_{1,1} f' + B_{1,0} f = 0\right]_{r=a} \text{ and } \left[B_{2,2} f'' + B_{2,1} f' + B_{2,0} f = 0\right]_{r=a},$$

with coefficients $B_{n,m}$ given below. As there is little evidence of buckling in the outer layer, and the Rachev-type contractility model generally yields tensile stresses that would inhibit buckling, we specify that the incremental deformation vanishes at the interface of the two layers with $f(b) = 0$ and $f'(b) = 0$ and that the incremental shear traction is zero (Yang et al., 2007), which gives

$$\left[C_{1,2} f'' + C_{2,1} f' = 0\right]_{r=b} \text{ and } \left[C_{2,2} f'' + C_{1,1} f = 0\right]_{r=b},$$

with coefficients $C_{n,m}$ given below.

We used the compound matrix method (Lindsay and Rooney, 1992, Haughton and Orr, 1997), a modification of the determinantal method commonly used for linear bifurcation analysis (Haughton and Ogden, 1979), to solve the differential equation numerically. The fourth order equation above was rewritten as a system of four, first order differential equations, $\boldsymbol{y}' = \boldsymbol{A}\boldsymbol{y}$, where $\boldsymbol{y} = [f, f', f'', f''']^T$ and $\boldsymbol{A}$ is the corresponding coefficient matrix. Boundary conditions at the inner surface and interface were similarly re-written as $[\boldsymbol{B}\boldsymbol{y} = \boldsymbol{0}]_a$ and $[\boldsymbol{C}\boldsymbol{y} = \boldsymbol{0}]_b$, respectively. We define two linearly independent initial conditions at $r = a$, which can be integrated to $r = b$ to create linearly independent solutions $\boldsymbol{y}^{(1)}$ and $\boldsymbol{y}^{(2)}$ such that $\boldsymbol{y} = k_1 \boldsymbol{y}^{(1)} + k_2 \boldsymbol{y}^{(2)} = \boldsymbol{M}\boldsymbol{k}$. For the determinantal method, one iterates on $\mathrm{T}_{act}$ until $\det([\boldsymbol{C}\boldsymbol{M}]_b) = 0$. However, this method can fail for stiff systems, thus we use Laplace expansions to write a new bifurcation condition equivalent to the original with $\det(\boldsymbol{C}\boldsymbol{M}) = \sum_k |C_k| \, \phi_k$ where

$$\phi_1 = \begin{vmatrix} y_1^{(1)} & y_1^{(2)} \\ y_2^{(1)} & y_2^{(2)} \end{vmatrix} = (1,2), \qquad \phi_2 = \begin{vmatrix} y_1^{(1)} & y_1^{(2)} \\ y_3^{(1)} & y_3^{(2)} \end{vmatrix} = (1,3),$$

and similarly, $\phi_3 = (1,4)$, $\phi_4 = (2,3)$, $\phi_5 = (2,4)$, $\phi_6 = (3,4)$. We evaluate $\mathcal{C}_k = |C_k|$ with

$$\mathcal{C}_1 = \begin{vmatrix} C_{1,1} & C_{2,1} \\ C_{1,2} & C_{2,2} \end{vmatrix} = (1,2)\ , \qquad \mathcal{C}_2 = \begin{vmatrix} C_{1,1} & C_{2,1} \\ C_{1,3} & C_{2,3} \end{vmatrix} = (1,3),$$

and similarly, $\mathcal{C}_3 = (1,4)$, $\mathcal{C}_4 = (2,3)$, $\mathcal{C}_5 = (2,4)$, $\mathcal{C}_6 = (3,4)$, where

$$(n,m) \equiv \begin{vmatrix} C_{1,n} & C_{2,n} \\ C_{1,m} & C_{2,m} \end{vmatrix}.$$

To evaluate the new bifurcation condition, we create a system of equations $\boldsymbol{\phi}' = \boldsymbol{\mathcal{A}}\boldsymbol{\phi}$ and identify the components of $\boldsymbol{\phi}'$ as, for example,

$$\phi_1' = \begin{vmatrix} y_1^{(1)\prime} & y_1^{(2)\prime} \\ y_2^{(1)} & y_2^{(2)} \end{vmatrix} + \begin{vmatrix} y_1^{(1)} & y_1^{(2)} \\ y_2^{(1)\prime} & y_2^{(2)\prime} \end{vmatrix} = \begin{vmatrix} \sum_i^4 A_{1i} y_i^{(1)} & \sum_i^4 A_{1i} y_i^{(2)} \\ y_2^{(1)} & y_2^{(2)} \end{vmatrix} + \begin{vmatrix} y_1^{(1)} & y_1^{(2)} \\ \sum_i^4 A_{2i} y_i^{(1)} & \sum_i^4 A_{2i} y_i^{(2)} \end{vmatrix}.$$

The components of $\boldsymbol{\mathcal{A}}$ can thus be conveniently defined in terms of the original components of $\boldsymbol{A}$ (Haughton and Orr, 1997), as listed below. This new system is then integrated from $a$ to $b$ using a fourth order Runge-Kutta method, and $\mathrm{T}_{act}$ is varied iteratively until the boundary condition at $b$ is satisfied, namely $[\sum_k \mathcal{C}_k\, \phi_k]_b = 0$. Loading conditions, including the volume change $v^*$, luminal pressure $P$, and axial stretch $\lambda_z$, can be varied parametrically to understand their effects on the critical value of $\mathrm{T}_{act}$ needed to induce buckling. Note, one may also fix the value of $\mathrm{T}_{act}$ and identify the critical value of a different loading variable of interest.

*Bifurcation Analysis – Specific Functions*

For the fourth order governing differential equation for the incremental displacement:

$$A_4 f'''' + A_3 f''' + A_2 f'' + A_1 f' + A_0 f = 0$$

we have

$$A_0 = (n^2 - 1)\left(r^2 \mathcal{B}''_{r\theta r\theta} + r\mathcal{B}'_{r\theta r\theta} + (n^2 - 1) * \mathcal{B}_{r\theta r\theta} + n^2(\mathcal{B}_{\theta\theta\theta\theta} - \mathcal{B}_{rrrr})\right)$$

$$A_1 = r\left(\left(r^2 \mathcal{B}''_{r\theta r\theta} + r\mathcal{B}'_{r\theta r\theta} - \mathcal{B}_{r\theta r\theta} - n^2(\mathcal{B}_{\theta\theta\theta\theta} + \mathcal{B}_{rrrr})\right) - n^2 r(\mathcal{B}'_{\theta\theta\theta\theta} + \mathcal{B}'_{rrrr})\right)$$

$A_2 = r^2(r^2\mathcal{B}''_{r\theta r\theta} + 7r\mathcal{B}'_{r\theta r\theta} + 5\mathcal{B}_{r\theta r\theta} - n^2(\mathcal{B}_{\theta\theta\theta\theta} + \mathcal{B}_{rrrr}))$

$A_3 = r^3(2r\mathcal{B}'_{r\theta r\theta} + 6\mathcal{B}_{r\theta r\theta})$

$A_4 = r^4\mathcal{B}_{r\theta r\theta}$

Note: These coefficients include only the non-zero components of $\boldsymbol{\mathcal{B}}$ for the considered stored energy density function.

For the fourth order differential equation

$$\boldsymbol{y}' = \boldsymbol{A}\boldsymbol{y} \text{ with } \boldsymbol{y} = [f, f', f'', f''']^T$$

note that

$$\boldsymbol{A} = \begin{bmatrix} 0 & 1 & 0 & 0 \\ 0 & 0 & 1 & 0 \\ 0 & 0 & 0 & 1 \\ -A_0/A_4 & -A_1/A_4 & -A_2/A_4 & -A_3/A_4 \end{bmatrix}.$$

For boundary conditions on the governing equation at the inner surface:

$$\left[B_{1,3}f''' + B_{1,2}f'' + B_{1,1}f' + B_{1,0}f = 0\right]_{r=a} \text{ and } \left[B_{2,2}f'' + B_{2,1}f' + B_{2,0}f = 0\right]_{r=a}$$

we have,

$B_{1,0} = (n^2 - 1)(r\mathcal{B}'_{r\theta r\theta} + \mathcal{B}_{r\theta r\theta})$

$B_{1,1} = r(r\mathcal{B}'_{r\theta r\theta} - (n^2 - 1)\mathcal{B}_{r\theta r\theta} + n^2(\mathcal{B}_{\theta\theta\theta\theta} + \mathcal{B}_{rrrr}))$

$B_{1,2} = r^2(r\mathcal{B}'_{r\theta r\theta} + 4\mathcal{B}_{r\theta r\theta})$

$B_{1,3} = r^3\mathcal{B}_{r\theta r\theta}$

$B_{2,0} = \mathcal{B}_{r\theta r\theta}(n^2 - 1)$

$B_{2,1} = \mathcal{B}_{r\theta r\theta}\, r$

$B_{2,2} = \mathcal{B}_{r\theta r\theta}\, r^2$

$B_{2,3} = 0$

For boundary conditions on the governing equation at the interface:

$$\left[C_{1,2}f'' + C_{1,1}f' = 0\right]_{r=b} \text{ and } \left[C_{2,2}f'' + C_{2,0}f = 0\right]_{r=b}$$

we have

$C_{1,0} = 0$

$C_{1,1} = \mathcal{B}_{r\theta r\theta}\, r$

$C_{1,2} = \mathcal{B}_{r\theta r\theta}\, r^2$

$C_{1,3} = 0$

$C_{2,0} = \mathcal{B}_{r\theta r\theta}\, (n^2 - 1)$

$C_{2,1} = 0$

$C_{2,2} = \mathcal{B}_{r\theta r\theta}\, r^2$

$C_{2,3} = 0$

Finally, for the compound matrix method component matrix

$$\boldsymbol{\mathcal{A}} = \begin{bmatrix} A_{11} + A_{22} & A_{23} & A_{24} & -A_{13} & -A_{14} & 0 \\ A_{32} & A_{11} + A_{33} & A_{34} & A_{12} & 0 & -A_{14} \\ A_{42} & A_{43} & A_{11} + A_{44} & 0 & A_{12} & A_{13} \\ -A_{31} & A_{21} & 0 & A_{22} + A_{33} & A_{34} & -A_{24} \\ -A_{41} & 0 & A_{21} & A_{43} & A_{22} + A_{44} & A_{23} \\ 0 & -A_{41} & A_{31} & -A_{42} & A_{32} & A_{33} + A_{44} \end{bmatrix}$$

with the components of $\boldsymbol{A}$ given above.

**Figure-S1**

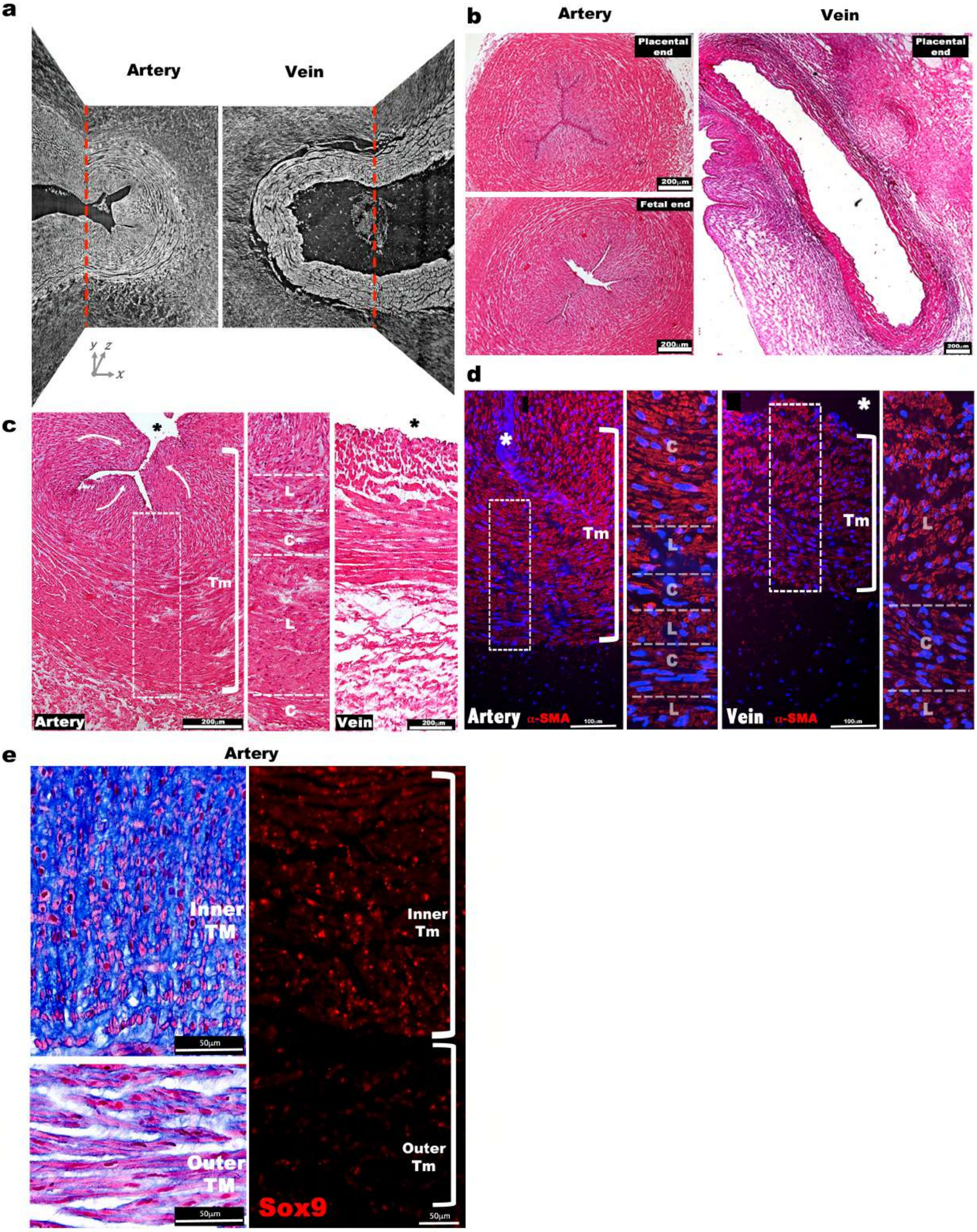

a
Artery
Vein
y z
x
b
Artery
Vein
Placental end
Fetal end
Placental end
200µm
200µm
200µm
c
Tm
L
C
L
C
Artery
200µm
Vein
200µm
d
Tm
C
L
C
L
C
L
Tm
L
C
L
Artery α-SMA
100µm
Vein α-SMA
100µm
e
Artery
Inner TM
50µm
Outer TM
50µm
Inner Tm
Outer Tm
Sox9
50µm

**Figure S1. Morphological and cellular characteristics of the human umbilical arteries and vein at birth.** (**a**) Synchrotron image from **Figure 1a** without red and purple shading illustrates contrast between inner and outer arterial tunica media as well as inner tuncia media buckling, compared with a uniform appearance of venous tunica media and lack of buckling. **(b)** Hematoxylin and eosin stained human umbilical artery cross-sections collected from the placental and fetal ends show occlusion of the umbilical artery with buckling of its interior, while the vein remains patent and lacks buckling. (**c**) Hematoxylin and eosin stained cross-sections show multiple cell layers composed of alternating circumferentially (C) or longitudinally (L) oriented smooth muscle cells (SMC) in the umbilical artery. White arrows indicate internal protrusion of the inner tunica media of the umbilical artery arising from buckling. (**d**) a-SMA (red) and DAPI (blue) staining of the umbilical artery and vein shows distinct orientation of the SMC layers as in **c**. (**e**) (*Left*) Alcian blue staining shows intense staining of the inner arterial tunica media (TM), with radially-oriented rounded cells contrasting with outer TM cells having typical SMC morphology. Eosin (pink) and nuclear fast red counterstaining on left. (*Right*) Sox9 immunostaining shows nuclear staining in SMCs of the inner umbilical artery TM . * marks the vessel lumen of in panels **c,d,e**. White brackets in **c,d** and **e** mark the TM. Tm, tunica media. Scale bars = 200μm in **b** and **c,** 100μm in **d** and 50μm in **e**.

### Figure-S2

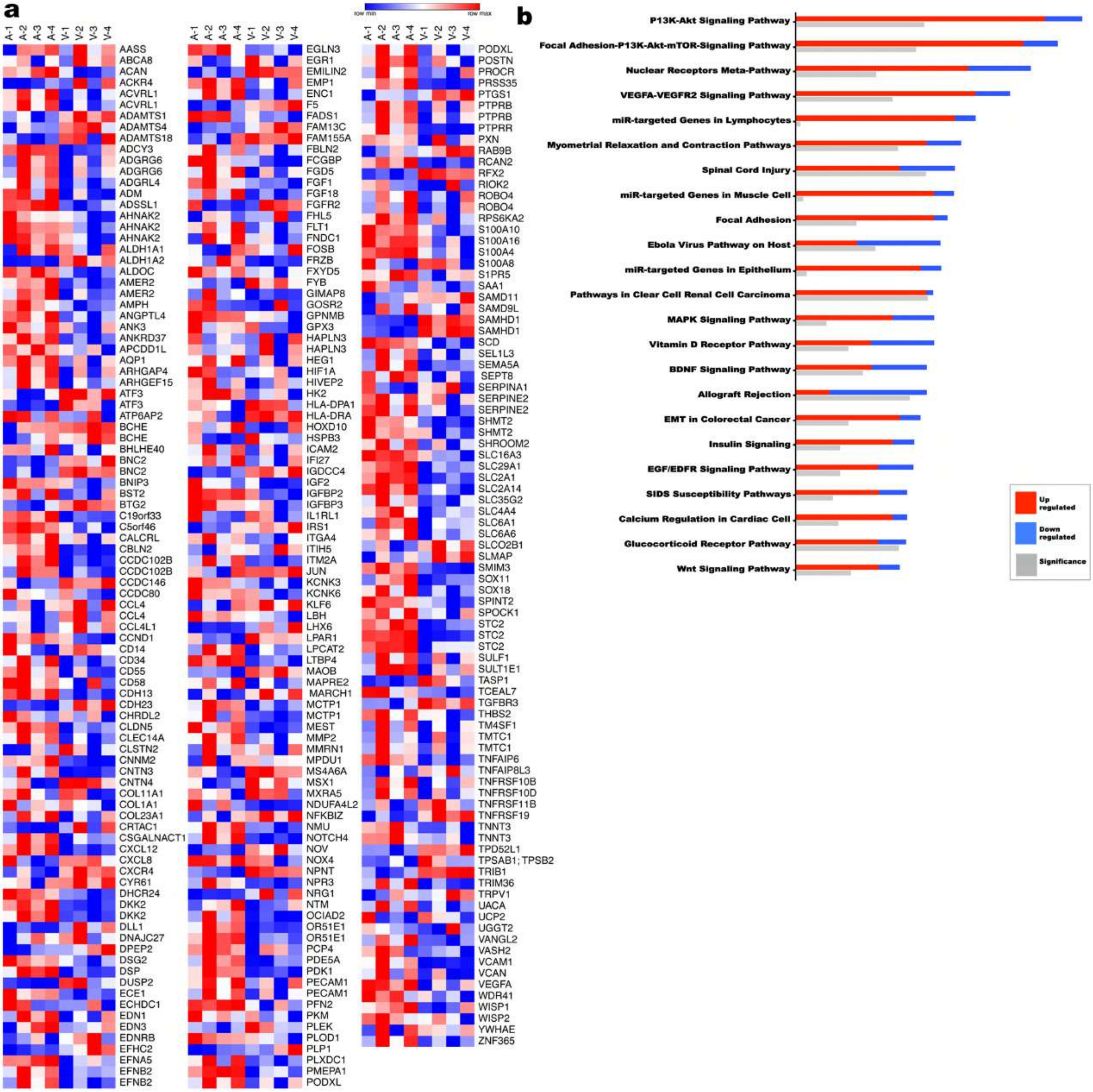


**Figure S2. Transcriptome comparison and pathway analysis of differences in the human umbilical artery and vein.** (**a**) Heat map of a subset of differentially expressed genes (1.9 fold change) from umbilical artery vs vein. (**b**) Ingenuity pathway analysis (IPA) summary of the most significantly different pathways.

### Figure-S3

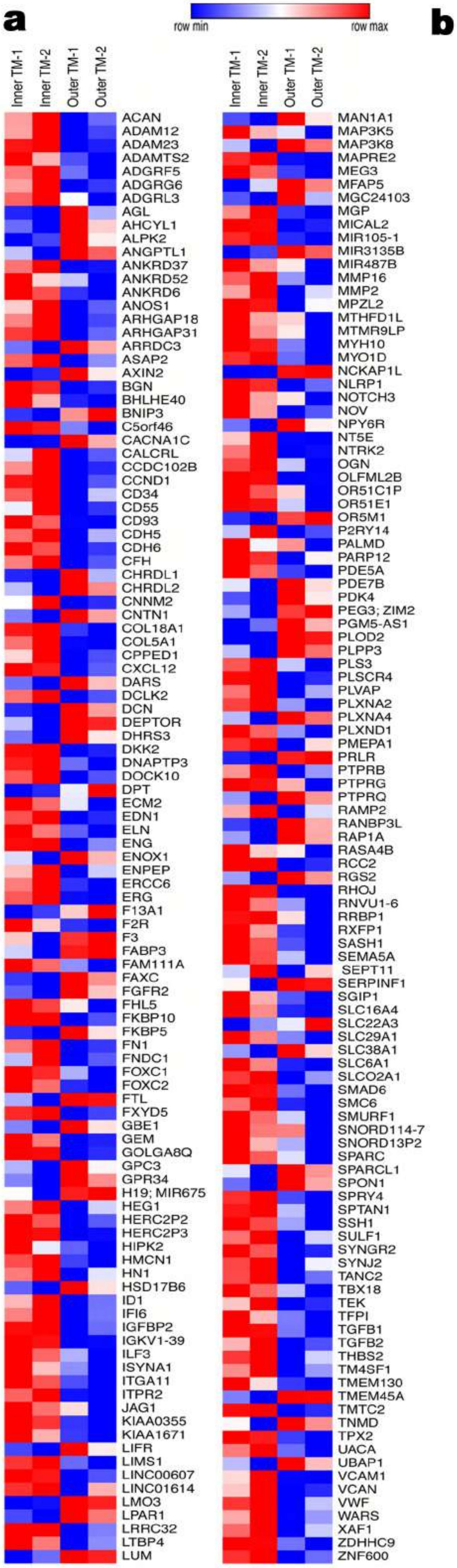


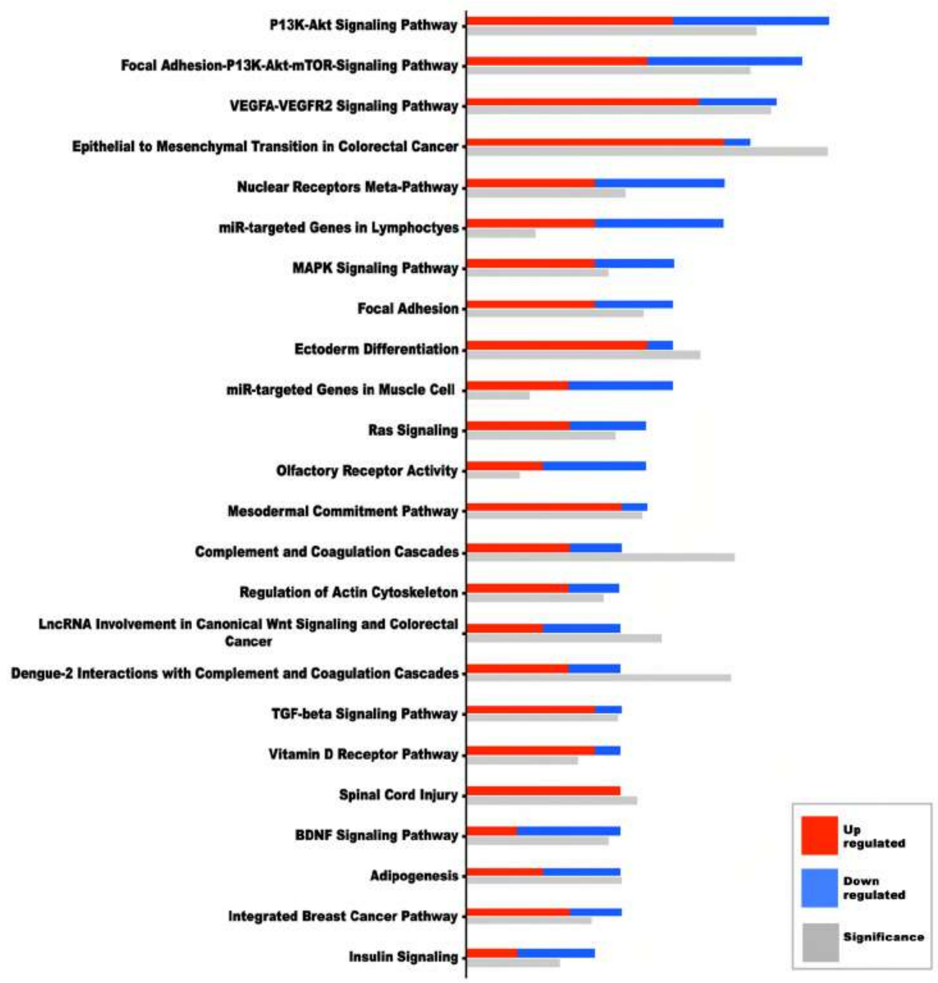


**Figure S3. Transcriptome comparison and pathway analysis of differences in the human inner umbilical artery tunica media vs the outer tunica media.** (**a**) Heat map of a subset of differentially expressed genes (1.9 fold change). (**b**) Ingenuity pathway analysis (IPA) showing significantly different pathways in the umbilical artery inner and outer tunica media.

**Figure-S4**

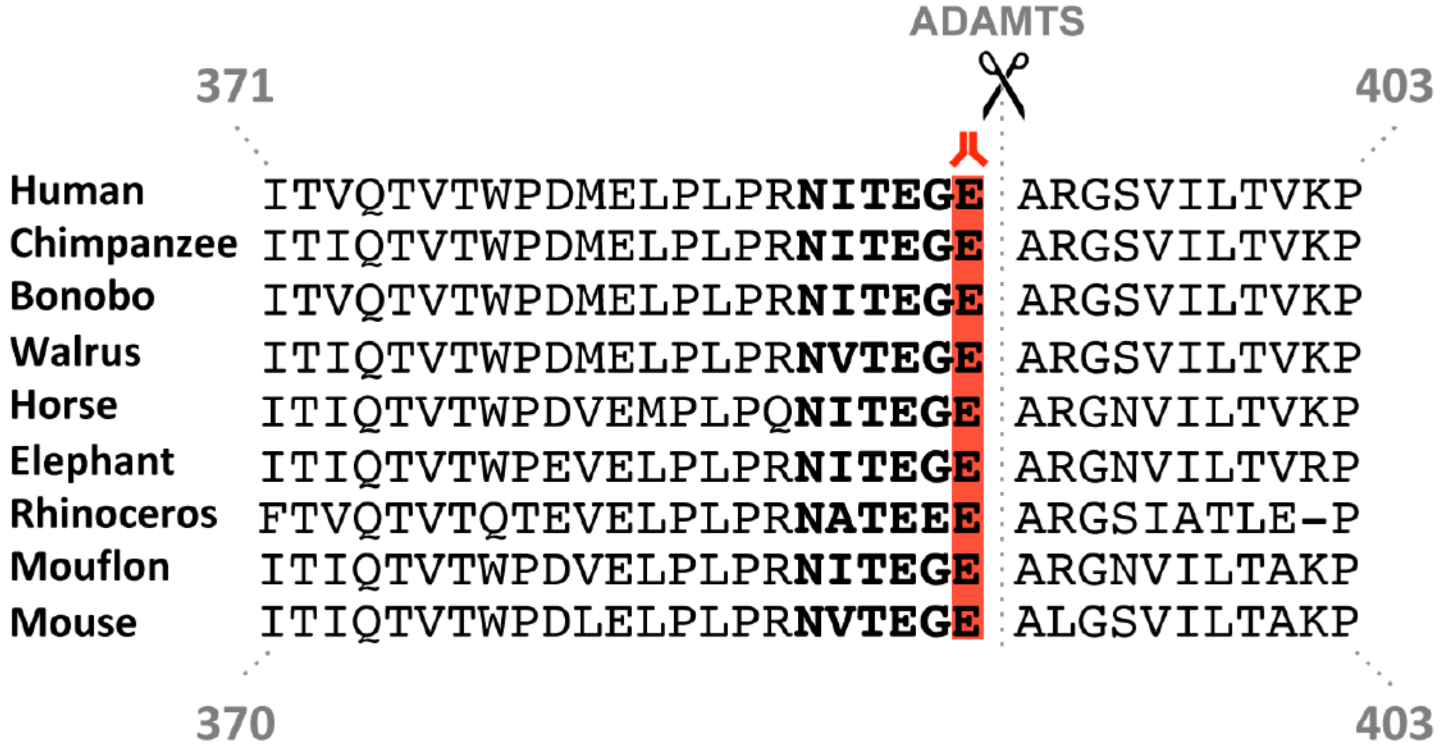


**Figure S4. Aggrecan cleavage site conservation in mammals.** Amino acid sequence spanning the ADAMTS-cleavage site in human aggrecan aligned with that of other mammalian species analyzed in this study. The anti-NITEGE antibody epitope (human) is highlighted in red.

**Figure-S5**

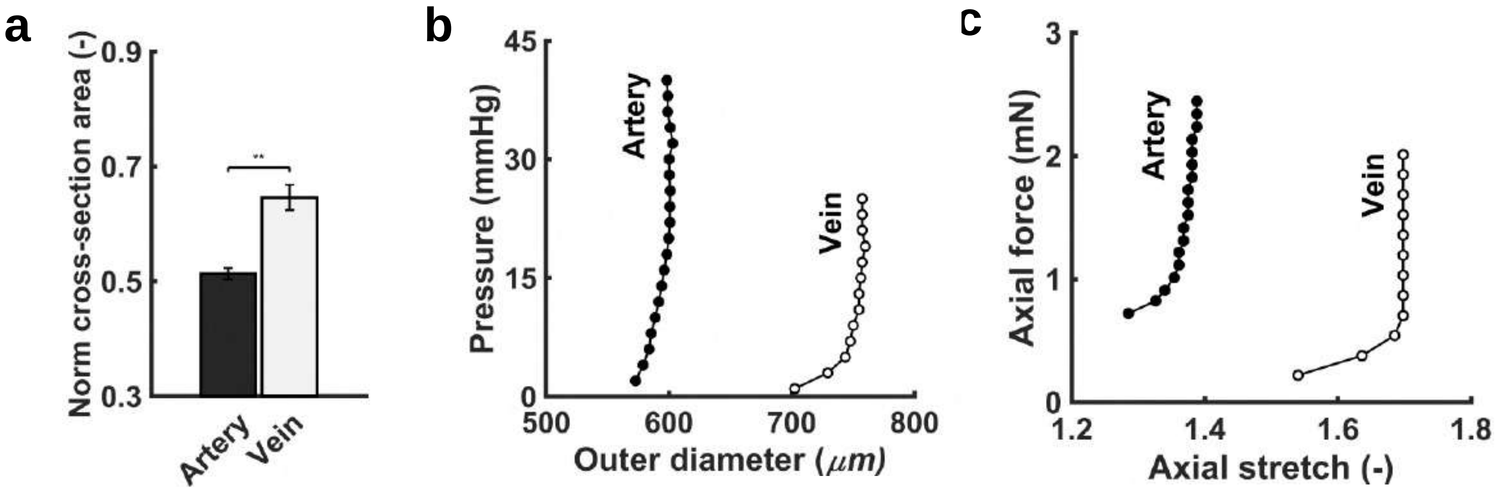


**Figure S5. Differential biomechanical properties of arteries and veins of the mouse umbilical cord. (a)** Shown are changes in cross-sectional area for the near term (E18.5) umbilical vein at a luminal pressure of 5 mmHg (vein) and the near term umbilical artery at 25 mmHg (artery) observed experimentally during isobaric contraction at vessel-specific fixed axial stretches. These data reveal a compressible behavior that aids in finally closure at birth **(b)** Nonlinear pressure-diameter behaviors during quasi-static pressurization at a fixed in vivo length for the umbilical artery and vein. Note the markedly different degrees of distension. **(c)** Nonlinear axial force-stretch behaviors during axial extension at a fixed luminal pressure for the umbilical artery and vein. Not the markedly different degrees of extension. Data from panels (b) and (c) thus show significant differences in the passive anisotropic behaviors, similar to the differential behaviors in the vasoconstrictive behaviors (Fig. 4c).

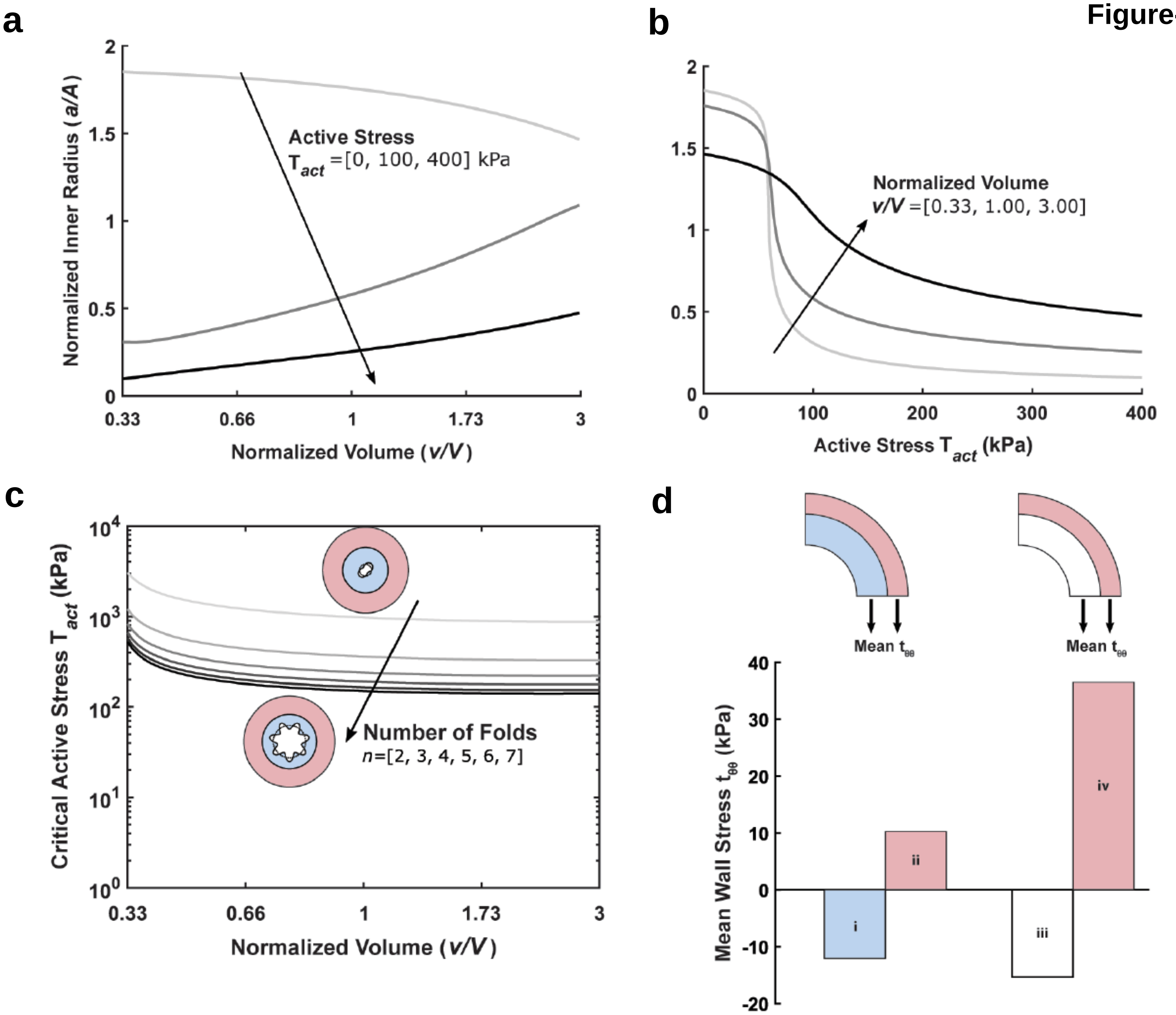


**Figure S6. Computational results for a model umbilical artery.** (**a**) Normalized inner radius as a function of normalized volume of the inner layer alone for different fixed values of the active stress parameter $T_{act}$ and (**b**) inner radius as a function of the active stress parameter $T_{act}$ for different fixed values of normalized volume in the inner layer. Loaded inner radius a was normalized by the unloaded inner radius A; the current volume of the vessel v was normalized by the original volume of the vessel V. **(c)** Critical value of the active stress parameter $T_{act}$ as a function of the normalized volume in the inner layer for different numbers of buckling-induced luminal folds n. **(d)** Mean circumferential stress $t_{\theta\theta}$ across the umbilical artery wall for varying normalized volumes: (i) and (ii) show circumferential stress for the case of shrinkage of the inner layer alone with v/V=0.5 while (iii) and (iv) show the case of no swelling with v/V=1.0. All simulations use the loading conditions, luminal pressure P=25 mmHg and fixed axial stretch $\lambda_z$=1.25.

**Figure-S7**

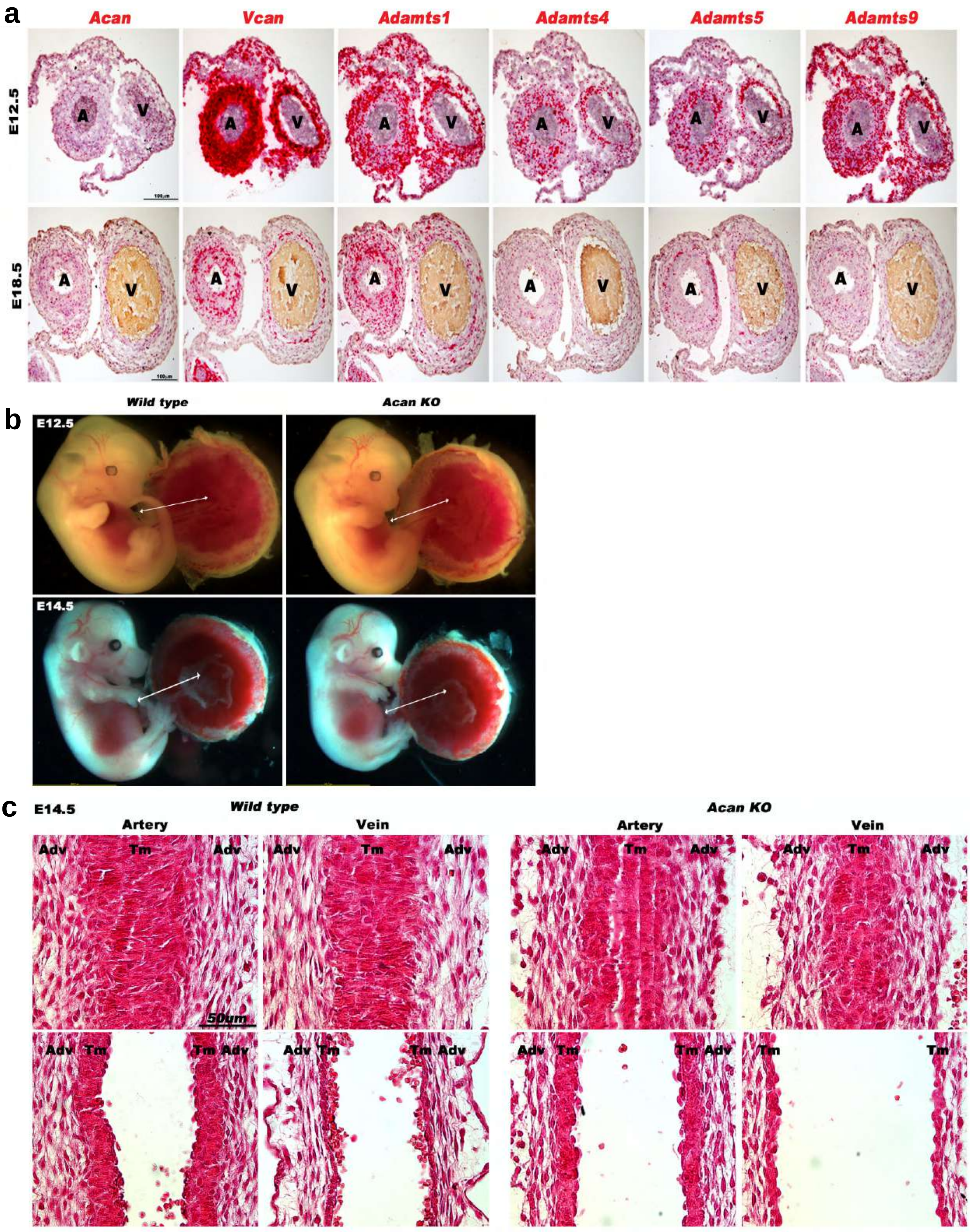
a
Acan
Vcan
Adamts1
Adamts4
Adamts5
Adamts9
E12.5
E18.5
A
V
100μm
b
Wild type
Acan KO
E12.5
E14.5
c
E14.5
Wild type
Acan KO
Artery
Vein
Adv
Tm
50um

**Figure S7. Gene expression and loss of aggrecan in the mouse umbilical cord.** (**a**) In situ hybridization analysis of E12.5 and E18.5 mouse umbilical cord sections for chondroitin sulfate proteoglycans and the ADAMTS metalloproteinases known to cleave them. (**b**) E12.5 and E14.5 wildtype and *Acan*$^{cmd/cmd}$ embryos have a comparable umbilical cord length. (**c**) Hematoxylin and eosin staining of E14.5 wild type and *Acan*$^{cmd/cmd}$ umbilical cords shows completion of longitudinal to circumferential reorientation of smooth muscle cells in the mutant arterial tunica media. Upper panels are from tangential sections through the arterial wall while the lower panels are taken through the approximate center of each vessel. Adv, adventitia; Tm, tunica media. Scale bars in **a**= 100μm and 50μm in **c.**
